# *Candida auris* evades innate immunity by using metabolic strategies to escape and kill macrophages while avoiding antimicrobial inflammation

**DOI:** 10.1101/2023.02.28.529319

**Authors:** Harshini Weerasinghe, Claudia Simm, Tirta Djajawi, Irma Tedja, Tricia L Lo, David Shasha, Naama Mizrahi, Françios AB Olivier, Mary Speir, Kate E. Lawlor, Ronen Ben-Ami, Ana Traven

## Abstract

*Candida auris* causes life-threatening, drug-resistant infections. In addition to drug resistance, therapeutic innovation is hindered by our limited knowledge of the mechanisms used by *C. auris* to evade immunity and establish infection. Here we show that *C. auris* escapes phagocytic containment and kills macrophages, and demonstrate that the mechanisms rely on metabolic regulation. We found that *C. auris*-infected macrophages undergo immunometabolic reprogramming and increase glycolysis but this does not lead to the expected antimicrobial responses, as macrophages fail to activate IL-1β cytokine and curb *C. auris* growth. Further analysis showed that *C. auris* relies on its own metabolic capacity to egress from macrophages, cause macrophage metabolic stress and cell death, and establish infection *in vivo*. We identified a transcriptional regulator of *C. auris* metabolism and macrophage evasion, and further show that, contrary to several other pathogens, *C. auris*-induced macrophage metabolic dysfunction and death fail to activate the NLRP3 inflammasome. Consequently, inflammasome-dependent antimicrobial responses remain inhibited throughout infection. Our findings establish a pivotal role for metabolic regulation in enabling *C. auris* to eliminate macrophages while remaining immunologically silent to ensure its own survival.

## INTRODUCTION

Drug-resistant microbial pathogens are one of the greatest threats to human health and medicine. The yeast *Candida auris* is one such pathogen, which emerged in clinical settings a little over a decade ago as a cause of bloodstream infections with high associated mortality (Jeffery-Smith et al., 2018; Nett, 2019). *C. auris* infections show intrinsic resistance to a front-line antifungal drug, common multidrug resistance and even pan-drug resistance in some cases, which is especially problematic given the small number of available antifungal therapies (Hanson et al., 2021; Jacobs et al., 2022; Jeffery-Smith et al., 2018; Nett, 2019).

In addition to antifungal drugs, the patient’s own innate immune system attempts to control fungal infections (Drummond et al., 2014). We are only beginning to understand how *C. auris* overcomes innate immunity to establish infection. Mannan in the *C. auris* cell wall has been implicated in hiding the immunostimulatory cell wall component ß-glucan (Wang et al., 2022). It has been suggested that mannan shields *C. auris* from recognition by macrophages and neutrophils, thereby reducing fungal killing and curbing pro-inflammatory cytokine production (Horton et al., 2021; Wang et al., 2022). However, another study concluded the opposite, showing that *C. auris* mannan induces more potent pro-inflammatory responses in human peripheral blood mononuclear cells (PBMCs) when compared to the common fungal pathogen *Candida albicans* (Bruno et al., 2020). Other than these contradictory studies on cell wall-mediated immune activation, no other mechanisms of immune evasion by *C. auris* have been identified.

In addition to modulation of cell surface antigens, immune evasion by microbial pathogens involves key metabolic adaptations to changing nutrient conditions in immune phagocytes (Traven and Naderer, 2019). Moreover, pathogen-induced changes in metabolic regulation and immunometabolism have emerged as major factors that regulate host immune responses and the ability of immune cells to control pathogens (Russell et al., 2019; Traven and Naderer, 2019; Troha and Ayres, 2020; Weerasinghe and Traven, 2020). Based on these studies, it has been proposed that metabolic interventions could promote beneficial immune, tissue and organ responses that protect host health during infection (Rosenberg et al., 2022; Traven and Naderer, 2019; Troha and Ayres, 2020). Clinical data backs this proposition, as maintaining metabolic homeostasis is critical in patients suffering from infections and sepsis (De Waele et al., 2020; Van Wyngene et al., 2018). As major reservoirs for the replication of antimicrobial resistant infections, macrophages have been studied for metabolic regulation during infection. Bacterial or fungal challenges cause a shift in macrophage metabolism that leads to increased glycolysis, reduced mitochondrial respiration and accumulation of TCA (tricarboxylic acid) cycle intermediates; collectively, these changes promote microbial killing and cytokine production (Cheng et al., 2016; Domínguez-Andrés et al., 2017; Gonçalves et al., 2020; Jha et al., 2015; Lachmandas et al., 2016; Michelucci et al., 2013; Mills et al., 2016; Tannahill et al., 2013). Pathogens can counter these host metabolic changes in several ways, for example by competitively using metabolites produced by macrophages for their own growth (Riquelme et al., 2020; Tomlinson et al., 2021) or modulating immune cell viability and function through nutrient competition (Tucey et al., 2018; Tucey et al., 2020; Wickersham et al., 2017). However, the metabolic mechanisms that control infection survival can vary dramatically between pathogens and therefore deep understanding is required across diverse infections (Traven and Naderer, 2019).

In this study, we establish key mechanisms of immune evasion by *C. auris*, which involve egress from immune containment by macrophages, killing of macrophages as well as evasion of recognition by a central macrophage immune regulator, the NLRP3 inflammasome. We show that these mechanisms depend on metabolic regulation in host and pathogen, identify a transcriptional regulator of metabolism in *C. auris* and show the importance of *C. auris* metabolism for macrophage killing and for establishing infection *in vivo*. Our study suggests that the metabolism-NLRP3-inflammation axis, a central response to microbial pathogens, is distinctly regulated with *C. auris*. Our findings further provide a framework for understanding how metabolism could be harnessed to modulate *C. auris* immune interactions that control infections.

## RESULTS

### *C. auris* escapes from macrophages

Very few studies have been published on the interaction between *C. auris* and macrophages. Therefore, to functionally define how *C. auris* impacts macrophage responses, we firstly assessed our *C. auris* strain for phagocytic uptake and fate in murine bone marrow-derived macrophages (BMDMs). Our *C. auris* isolate 470121 belongs to Clade I and was obtained from the Australian National Mycology Reference Centre.

*C. auris* was phagocytosed by BMDMs, with an average of 3.17 yeast per macrophage counted 1-hour post challenge with an MOI of 3 (Dataset S1). The number of yeast cells/macrophage exceeding the MOI could be due to yeast replication, as *C. auris* survived and replicated inside macrophages (Figure 1A). This result is consistent with our previous work with a different *C. auris* isolate and the work of Bruno et al (Bruno et al., 2020; Simm et al., 2022). Therefore, replication in macrophages is a fairly general capability of *C. auris* strains. At 8-10 hours post challenge *C. auris* escaped from macrophages, as evident by replication of yeast cells in the surrounding medium and a sharp increase in extracellular colony forming units (CFUs) (Figure 1A and 1B, Figure S1A and Movie S1). Notably, several escape events were observed from individual macrophages showing the efficiency of *C. auris* dissemination (Movie S2). There was no morphological change associated with the escape, as *C. auris* remained in yeast form throughout the experiment (Figure 1B, Movie S1 and S2). Replication within macrophages and *C. auris* escape was not limited to glucose but occurred in several carbon sources, such as galactose and mannose (Figure S1B). Differences in CFUs between carbon sources was attributed to the differing abilities of our *C. auris* strain to utilize them: *C. auris* growth was faster in glucose and mannose compared to media containing galactose or lacking in glucose (Figure S1C).

**Figure 1.**
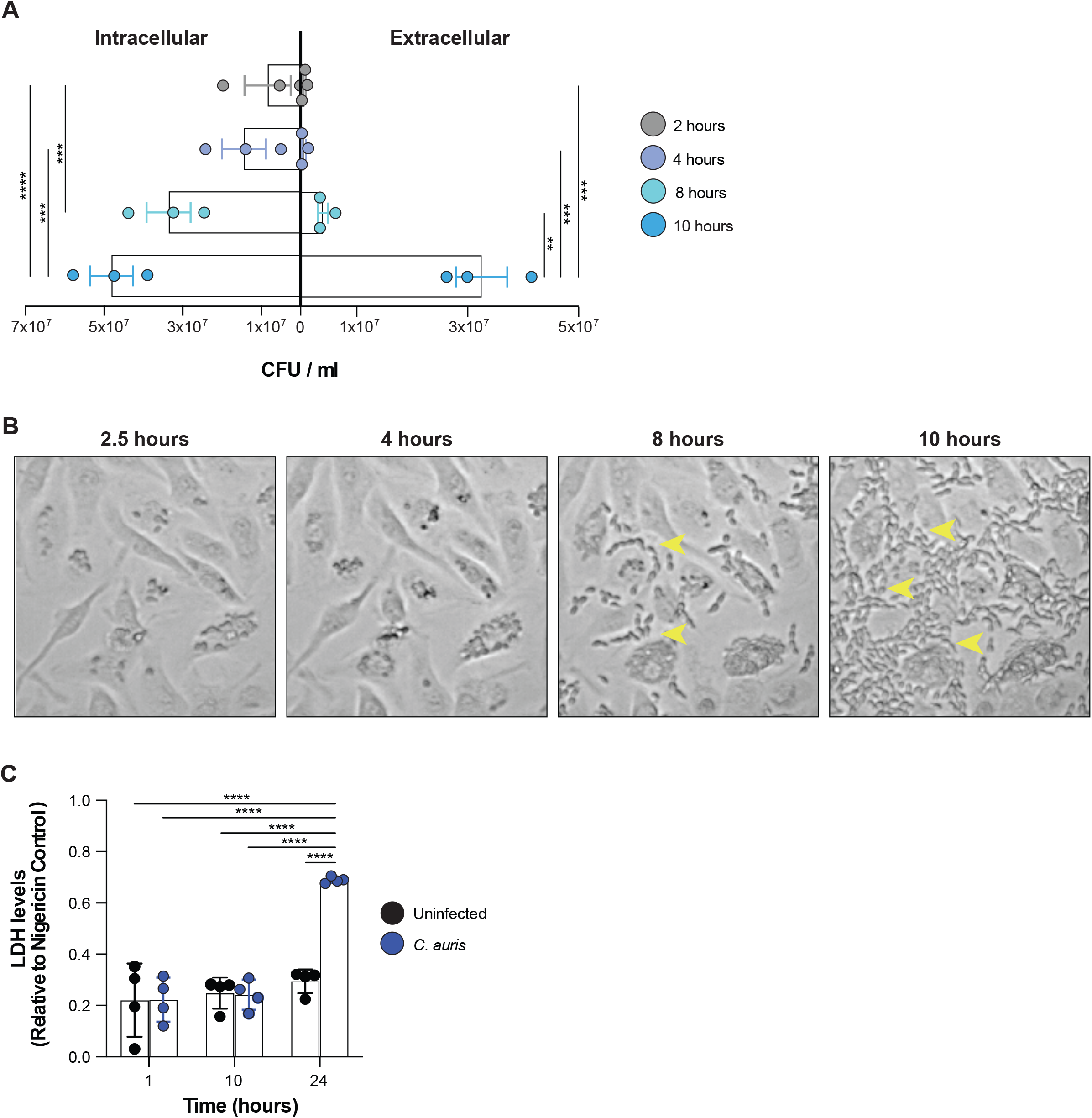
*C. auris* escapes from macrophages. **A.** Quantitation of colony forming units (CFUs) of *C. auris* 470121 for intracellular growth in macrophages (BMDMs) and extracellular growth in the surrounding medium. MOI was 3:1 (*C. auris*: macrophages). Supernatants and macrophage lysates were plated at 2, 4, 8 and 10 hours post challenge and CFUs counted after 2 days of incubation at 30 °C. The bar graph shows mean values and SEM from 3 independent experiments. Statistical significance was determined using the 2-way ANOVA Bonferroni’s multiple comparison test (***p≤0.001, **** p≤0.0001). Only statistically significant comparisons are indicated in the figure. **B.** Still microscopy images from Movie S1 of *C. auris* infecting and escaping mouse BMDMs at 2, 4, 8 and 10 hours post challenge. Images are magnifications of a section of the field of view (the entire field of view and the magnified sections are shown in Figure S1B). Yellow arrowheads indicate yeast cells of *C. auris* that have escaped macrophages (8 and 10 hour time point). Movie S2 shows escape events as capture by live cell imaging in individual macrophages. **C.** The release of lactate dehydrogenase (LDH) into the supernatant. Measurements were performed for uninfected and *C. auris-*infected macrophages at 1, 10 and 24 hours post challenge. Treatment with the NLRP3 inflammasome activator nigericin causes wide-spread macrophage lysis and was used as a control. All values were expressed relative to LDH release from nigericin-treated macrophages. The bars represent the mean values and SEM from 4 independent experiments. A 2-way ANOVA Bonferroni’s multiple comparison test was performed (**** p≤0.0001). Only statistically significant comparisons are indicated in the figure.

Intriguingly, despite abundant extracellular yeast growth being evident at 10 hours post *C. auris* infection (Figure 1A), there was no associated increase in lactate dehydrogenase (LDH) release as a measure of macrophage plasma membrane damage (Figure 1C). Therefore, *C. auris* is able to effectively egress from macrophages to avoid immune containment and promote dissemination, but this is not associated with wide-spread macrophage lysis.

### *C. auris* kills macrophages

Extending the assay time course showed that, at 24 hours post infection, LDH release increased significantly from *C. auris*-infected macrophages (Figure 1C). This result indicated that, although the initial yeast escape does not trigger macrophage lysis, at later time points in infection *C. auris* causes lytic immune cell death. To further understand the dynamics of macrophage killing by *C. auris*, we employed our live-cell imaging platform that uses nuclear staining with the dye DRAQ7 as a measure of macrophage membrane permeabilization and cell death, and quantifies these events at single cell level in large macrophage populations (Tucey et al., 2016). Live cell imaging confirmed that macrophages survived the initial escape of *C. auris* and did not succumb for several hours afterwards (Figure 2A and 2B, Movie S1). Rapid macrophage membrane permeabilization initiated at around 16-18 hours after challenge. It reached almost 100% of DRAQ7-positive macrophages by 24 hours with a MOI of 6 and close to 70% when MOI of 3 was used (Figure 2A). Thus, *C. auris* kills macrophages effectively and is able to eliminate the entire macrophage population. The escape and killing of mouse macrophages by *C. auris* (Figure 2A and 2B) was fully recapitulated in human monocyte-derived macrophages (hMDMs) and was triggered by active *C. auris*, as treatment with the antifungal drug caspofungin inhibited macrophage cell death (Figure 2C).

**Figure 2.**
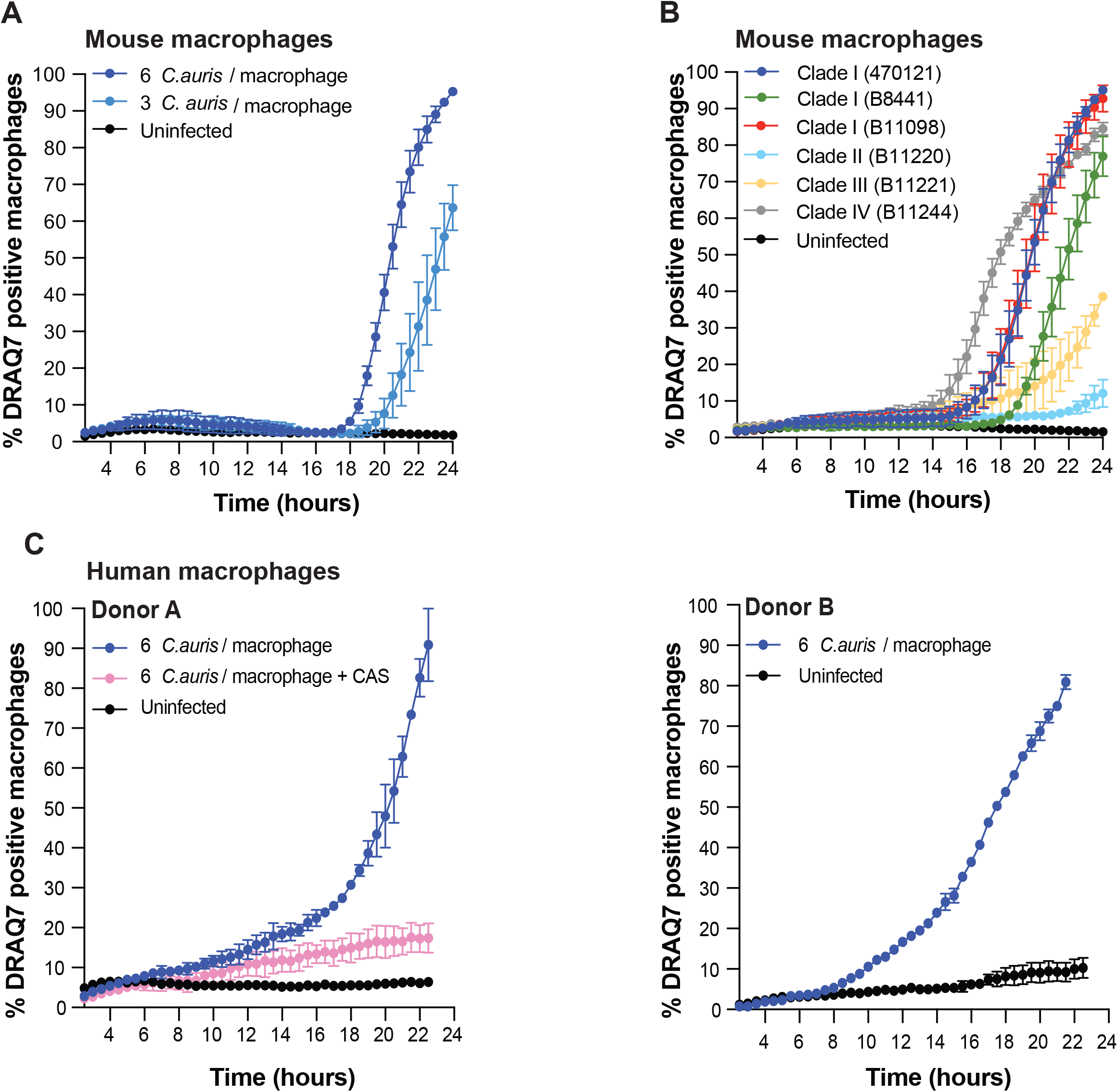
*C. auris* kills mouse and human macrophages. **A.** Macrophages (BMDMs) were challenged with *C. auris* strain 470121 (MOI 3 or 6) and imaged over 24 hours. Macrophage membrane permeabilization was determined using the membrane-impermeable dye DRAQ7 and DRAQ7-positive nuclei counted over time in macrophage populations. Shown are the mean and SEM for 3 independent experiments. **B.** Same as A, but human monocyte-derived macrophages (hMDMs) were used to determine if killing of mouse macrophages by *C. auris* is conserved in human macrophages. *C. auris* strain 470121 was used for these experiments, with hMDMs isolated from two independent healthy donors (donor A and donor B). MOI was 6. For Donor A caspofungin was added to the media at the concentration of 125 ng/ml. Donor A and B each represent two independent experiments consisting of two technical repeats for every condition and shown here are the mean values and SEM values for these. **C.** Same as A, but BMDM challenge was performed using a compendium of *C. auris* isolates belonging to clades I-IV as indicated. The MOI was 6. Shown here are the mean values and SEM for 3 independent experiments.

We wanted to further ascertain if the ability to escape and kill macrophages was conserved between diverse clinical isolates belonging to several genomic clades (Lockhart et al., 2017). *C. auris* strains from clades I-IV were obtained from the CDC and used in time-lapse imaging assays quantifying macrophage viability. The escape and killing of macrophages by *C. auris* were conserved across clades I-IV (Figure 2B). There were however differences in the kinetics. The clade IV isolate was the fastest in triggering macrophage cell death, while the clade II and clade III isolates were the slowest (Figure 2B). The clade IV strain B11244 triggered macrophage cell death two hours earlier than all other clades (starting at 14-16 hours), while the clade II strain B11220 and the clade III strain B11241 showed a delay of two or four hours respectively (Figure 2B). Some of these differences can be explained by the growth rates of the isolates. Clade II and clade III isolates were the slowest to kill macrophages and were also the slowest growing isolates *in vitro* (Figure S2A). However, growth rates do not explain all of the differences because the clade IV strain B11244 was the fastest in triggering macrophage cell death, but it was not the fastest growing strain of the group (Figure S2A).

### *C. auris* kills macrophages by causing metabolic stress

The findings in Figure 2 suggested that active proliferation of *C. auris* was important for inducing macrophage death. We hypothesised two possible mechanisms: (i) *C. auris* is competing for nutrients required to sustain macrophage viability or (ii) *C. auris* is secreting a compound toxic to macrophages.

Central carbon metabolism is critical for infected macrophages, reviewed in (Traven and Naderer, 2019). Therefore, to test the nutrient competition hypothesis, we supplemented macrophage growth media with metabolites that feed into central carbon metabolism (Figure 3A) and then asked if they were able to rescue macrophages from *C. auris*-induced cell death. Supplementation of glucose or pyruvate delayed death of *C. auris*-infected macrophages, while supplementation of arginine, glutamine or acetate did not (Figure 3B). To rule out osmotic effects from additional levels of nutrients delaying membrane lysis and cell death, we supplemented sorbitol at the same concentration (10 mM). Sorbitol did not rescue macrophage viability upon *C. auris* infection (Figure 3B). Taken together, these data show that glycolysis is essential for *C. auris*-infected macrophages to make pyruvate feeding into the TCA cycle, thereby producing the energy required to enhance survival.

**Figure 3.**
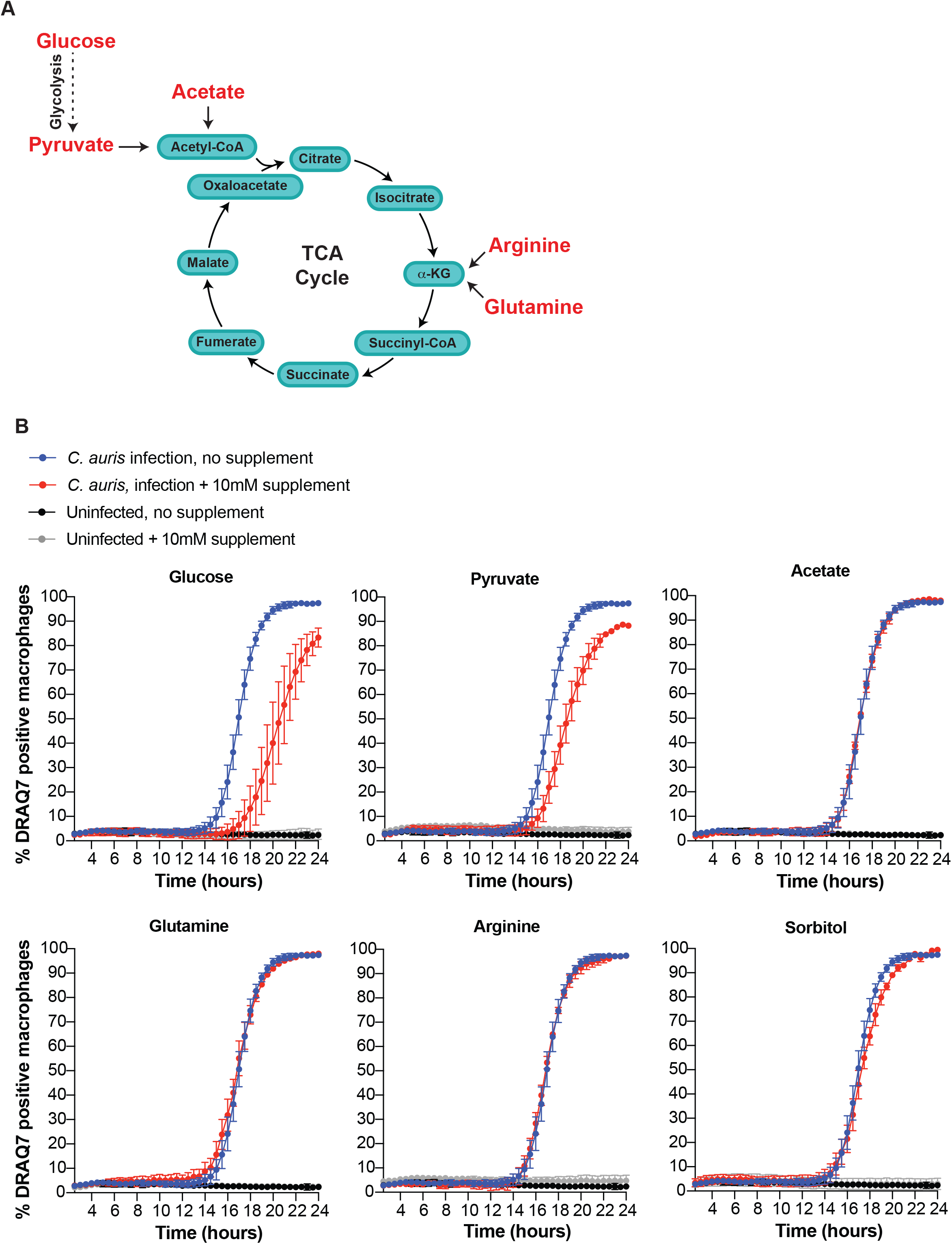
*C. auris*-infected macrophages can be rescued by nutrient supplementation. **A.** Central carbon metabolism (glycolysis and the TCA cycle). The nutrients indicated in red were supplemented to the medium in *C. auris*-macrophage infection conditions to test if they can rescue macrophages from *C. aur*is-induced cell death. **B.** Macrophages (BMDMs) were challenged with *C. auris* (strain 470121) and DRAQ7-positive macrophages determined by live cell imaging over 24 h. The indicated nutrients were added to standard macrophage infection medium which contains 10 mM glucose. The blue graphs (labelled as *C. auris*) are from infections performed in standard medium without supplementation of additional nutrients. All of the nutrient conditions were analysed together in the same experiment but are plotted separately for clarity. As such, the standard medium (blue graph) is the same in all comparisons. Shown here are the mean values and SEM for n=3 independent experiments.

These results were surprising in light of previous work that suggested that infection with *C. auris* does not trigger a metabolic shift in PBMCs towards higher glucose consumption (Bruno et al., 2020). We therefore further assessed the metabolism of macrophages in response to *C. auris* by studying their metabolic gene signatures. *C. auris* infection induced the mRNA expression of the *Glut1* glucose transporter and the glycolytic enzymes *Hk2* and *Pfkbf3* (Figure 4A), as is characteristic when immune cells increase glycolysis (Freemerman et al., 2014; Kelly and O’Neill, 2015; Rodríguez-Prados et al., 2010; Tucey et al., 2018). Infected macrophages also showed further signs of elevated glycolysis with heightened levels of lactate relative to uninfected controls (Figure 4B) and accelerated cell death in the presence of the mitochondrial complex I inhibitor metformin (Figure 4C). At the dose used, metformin was not toxic to uninfected macrophages (Figure 4C). Hyper-susceptibility to metformin is in line with *C. auris* triggering the Warburg effect in macrophages. During Warburg metabolism, decreased mitochondrial respiration accompanies higher glycolysis, which could make glucose essential for macrophage survival. Indeed, depletion of glucose was observed as *C. auris* infection progressed (Figure 4D) and supplementation of glucose rescued *C. auris*-infected macrophages in a dose-dependent manner (Figure 4E). The viability of uninfected macrophages did not change dependent on glucose concentrations and was maintained in medium with no glucose (Figure 4E).

**Figure 4.**
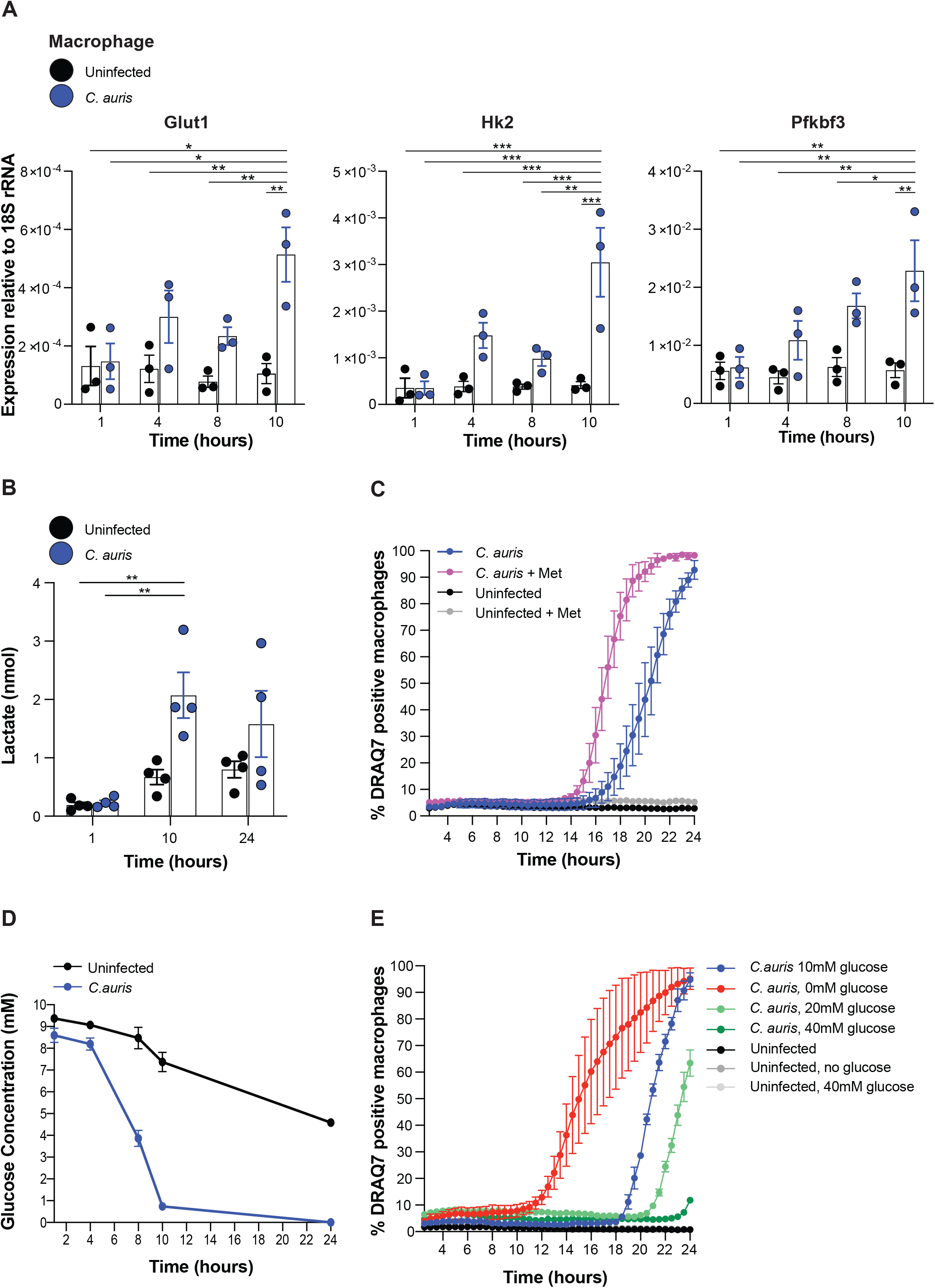
Immunometabolic adaptation of macrophages in response to *C. auris*. For all experiments, *C. auris* challenge was performed with BMDMs. The fungal strain was 470121 and the MOI was 6 unless otherwise stated. **A.** Quantitative PCR analysis of the expression of macrophage metabolic genes: the glucose importer *Glut1* and the glycolytic enzyme genes *Pfkfb3* and *Hk2* at 1, 4, 8 and 10 hours after challenge with *C. auris*. Gene expression was normalised to 18S rRNA. Shown are the mean values and SEM for 3 independent experiments, each analysed in two technical replicates. A 2-way ANOVA Bonferroni’s multiple comparison test was performed (* p≤0.05, ** p≤0.01, *** p≤0.001,). Only statistically significant comparisons are indicated in the figure. **B.** Production of lactate was determined for *C. auris-*infected and uninfected BMDMs 1, 10 and 24 hours after challenge. Shown are the means and SEM from 4 independent experiments. A 2-way ANOVA Bonferroni’s multiple comparison test was performed to compare the 10 and 24-hours to the 1-hour infection conditions (** p≤0.01). Only statistically significant comparisons are indicated in the figure. **C.** Effects of metformin on *C. auris*-infected macrophages. Macrophages were pre-treated with 5mM metformin for 24 hours and then switched to fresh media with and without metformin, followed by challenge with *C. auris*. Uninfected macrophage controls were treated the same. Shown here are the mean values and SEM for 3 independent experiments. **D.** Reduction in glucose levels over the time course of *C. auris-*macrophage infection. Glucose measurements were taken from equal number of both infected and uninfected macrophage conditions. Shown here are the mean values and SEM for n=3 independent experiments. **E.** Macrophage cell death during *C. auris* infection in media containing increasing concentrations of glucose. The control condition (blue) contains 10 mM glucose. Shown here are the mean values and SEM for n=3 independent experiments.

Increasing glucose concentrations also led to increased growth of *C. auris* (Figure S2B). This means that, although in higher glucose *C. auris* was proliferating more robustly, macrophages nevertheless survived better. These data argue against the hypothesis that *C. auris* could be secreting metabolites that are toxic to macrophages, as these toxic metabolites would be expected to increase when yeast growth is more robust in higher glucose, yet we observed less macrophage cell death under these conditions. Collectively, our results show that macrophages increase their glycolytic metabolism in response to *C. auris* but are unable to inhibit *C. auris* growth or kill the pathogen. We further show that the mechanism by which *C. auris* kills macrophage is explained by glucose competition with macrophages, which causes host metabolic stress by depleting glucose and disabling host glycolysis.

### *C. auris* metabolic capacity drives macrophage escape and killing and is important for establishing infection *in vivo*

To understand how *C. auris* utilises and regulates its metabolic capacity during macrophage interactions, we created a mutant unable to perform glycolysis by deleting the gene encoding pyruvate kinase (*CDC19*). As expected, the *Δcdc19* mutant was unable to utilise glucose or fructose as a carbon source but grew on acetate albeit more slowly than the wild type (Figure 5A and Figure S3A). The *Δcdc19* mutant was unable to either escape or kill macrophages, even when acetate was supplemented to the medium to enable its growth (Figure 5B). This supports a key role of *C. auris* metabolic capacity in both immune egress and killing.

**Figure 5.**
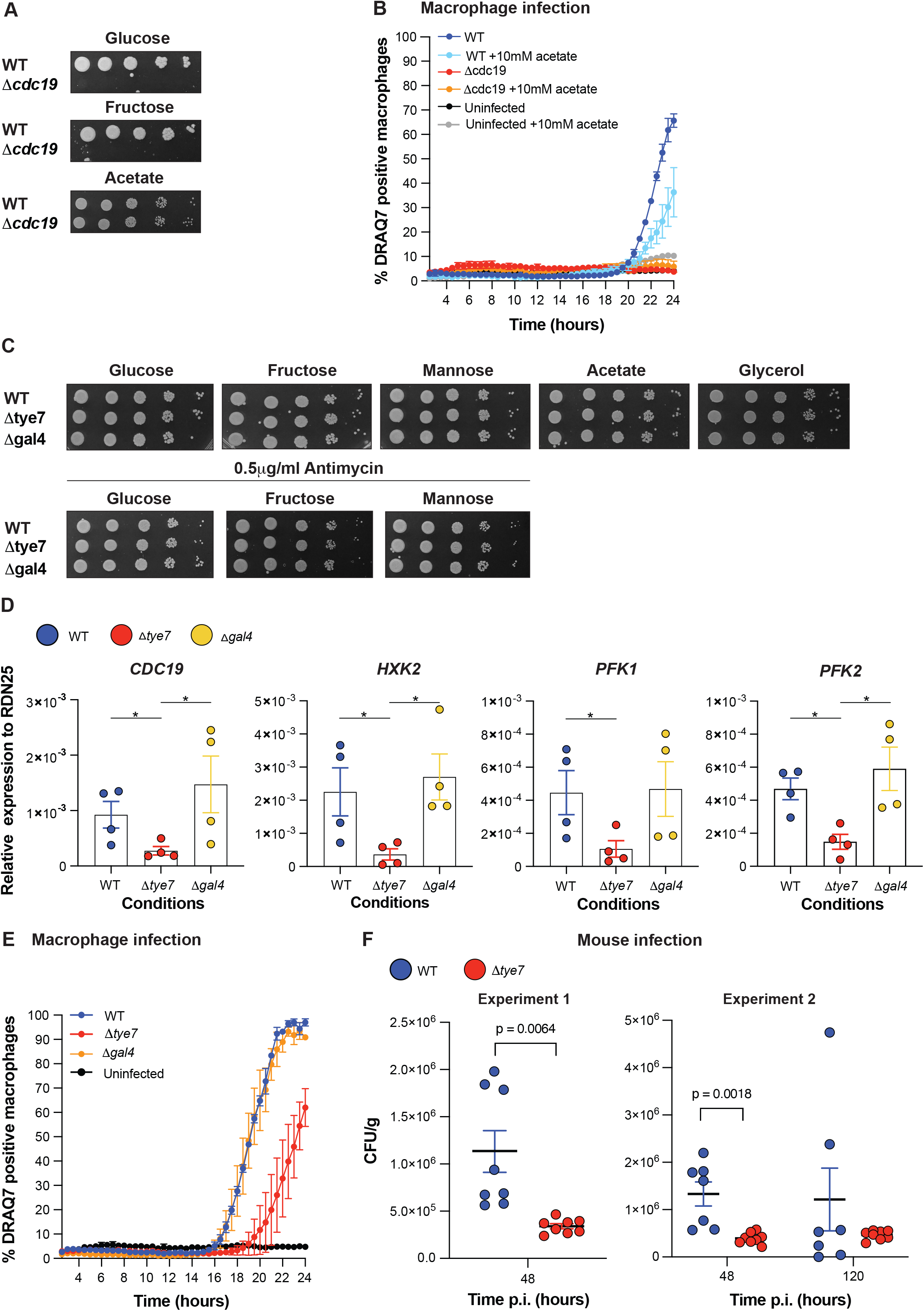
Roles of *C. auris* metabolic capacity in macrophage killing and establishment of *in vivo* infection in organs. **A.** Spot assays of the *C. auris* pyruvate kinase mutant *Δcdc19* relative to its isogenic wild type (strain B8441) in media containing glucose, fructose or acetate as the carbon source. Growth was assessed after 3 days at 30 °C. **B.** Macrophages (BMDMs) were challenged with *C. auris* wild type (strain B8441) or its isogenic *Δcdc19* strain (MOI 6). The uninfected controls are also included. DRAQ7-positive macrophages were quantified using live-cell imaging. Graphs show the mean values and SEM for n=3 independent experiments. **C.** Spot assays for wild type, *Δtye7* and *Δgal4* mutants of *C. auris*. Cultures were plated on the indicated plates and photographed after 3 days of growth at 30 °C. **D.** Yeast cultures were grown as in Figure S3B and samples taken after 7 hours, when both wild type and mutants enter log phase. The expression of the indicated glycolysis genes was assayed using qPCR and normalised to *RDN25*. Shown are the means and SEM of n=4 independent experiments. *p≤0.05 (Kruskal Wallis non-parametric test and the Dunn’s multi-comparison test). **E.** Macrophages (BMDMs) were challenged with *C. auris* wild type (470121), *Δtye7* and *Δgal4* and DRAQ7-positive macrophages quantified using live-cell imaging. The MOI was 6. Shown are the mean values and SEM for n=3 independent experiments. **F.** Mice were infected with *C. auris* wild type or *Δtye7* (470121 background) and kidney CFUs determined after 48 hours. Two different experiments were performed. In the second experiment, a further time point of 120 hours was sampled. Statistical analysis was performed using the Kruskal Wallis non-parametric test and the Dunn’s multi-comparison test.

We next aimed to understand how *C. auris* regulates its metabolic capacity to kill macrophages and establish infection. For this, we created mutants in the transcriptional regulators Tye7 and Gal4. Tye7 is a transcriptional activator that controls glycolysis in yeasts such as *C. albicans* and the distantly related model organism *S. cerevisiae* (Askew et al., 2009; Nishi et al., 1995). In *C. albicans* the activator Gal4 further contributes to glycolytic gene expression (Askew et al., 2009), although *S. cerevisiae* Gal4 has a different function in regulating galactose utilisation (Traven et al., 2006). The *C. auris Δtye7* and *Δgal4* mutants grew in all carbon sources that we tested (fermentable and non-fermentable), even in the presence of the mitochondrial respiration inhibitor antimycin A that forces cells to use glycolysis (Figure 5C). A more detailed analyses of growth and doubling time in liquid medium confirmed that *Δgal4* displayed wild type growth rates in glucose, which was consistent in two different strain backgrounds (470121 and B8441) (Figure S3B and S3C). This analysis revealed that *Δtye7* grew more slowly in log phase, with the average doubling time of almost 2 or 1.5 times the rate of the wild type for the 470121 and B8441 strains respectively (Figure S3B-S3E). However, *Δtye7* entered log phase normally and reached the same saturation as the wild type strain at the conclusion of the time course (Figure S3B and S3C).

To test whether these transcription factors regulate glycolysis in *C. auris*, we firstly examined the expression of relevant glycolytic genes. Fungal cultures were grown as for the growth curves in Figure S3B, and samples taken at the 7-hour time point when both wild type and mutants entered log phase and before any difference in growth rate occurs. The expression of glycolytic genes was reduced in *Δtye7* but not in *Δgal4* (Figure 5D). This suggests that Tye7 is a prominent transcriptional activator of glycolytic gene expression in *C. auris*. Consistent with this key metabolic role, the *Δtye7* mutant was slower than the wild type in triggering glucose-dependent macrophage cell death (Figure 5E). The *Δgal4* mutant did not differ from the wild type (Figure 5E). Crucially, glycolytic metabolism and metabolic adaptation were also important to establish infection *in vivo*, as shown by reduced *C. auris* proliferation in the kidneys of mice systemically infected with the *Δtye7* strain at day 2 after infection (Figure 5F). Given the slower growth of the *Δtye7* strain *in vitro* (Figure S3), we extended the post infection time to 5 days to address if the *Δtye7* strain simply needs more time to proliferate in mouse kidneys. However, CFU numbers for *Δtye7* did not increase at day 5 (Figure 5F), showing that even after prolonged times the mutant is unable to grow in kidney tissues. By day 5, the wild type CFUs decreased (Figure 5F), indicating that mice were clearing wild type infection. Collectively, these results show that the metabolic capacity of *C. auris* and its glycolytic metabolism are important for evading macrophages and for establishing infection in host organs. Our data further identified Tye7 as a key transcriptional regulator metabolism in *C. auris*.

### *C. auris* avoids recognition by the NLRP3 inflammasome

Next, we tested if *C. auris* pays the price for causing macrophage metabolic stress and cell death. We tested this because macrophage cells death is known to alert the host to infection and trigger inflammation (Traven and Naderer, 2014). Moreover, it has been shown that macrophages can sense microbial metabolism, because disruption of macrophage glycolysis by infection activates the NLRP3 inflammasome in response to several pathogens (Sanman et al., 2016; Tucey et al., 2020; Wolf et al., 2016). This then serves to control infection, as NLRP3 inflammasome triggers antimicrobial inflammation *via* maturation of the proinflammatory cytokines IL-1β and IL-18, as well as activating pyroptosis, a proinflammatory type of macrophage cell death that involves caspase-1/4/5/11-mediated cleavage and activation of the pore forming protein Gasdermin D (GSDMD) (Chen and Schroder, 2013; Yabal et al., 2019).

We therefore wondered if, as is the case for other pathogens, *C. auris*-induced metabolic stress and cell death of macrophages alert the host to infection, leading to NLRP3-inflammasome dependent antimicrobial inflammation. However, this was not the case for our *C. auris* isolate. By ELISA, we detected little bioactive IL-1β in the supernatants of *C. auris*-infected macrophages at 1 hour after challenge (Figure 6A). Secreted IL-1β increased at 10 hours, but the concentration was low relative to the nigericin control and there was no additional increase at 16 hours (Figure 6A). Measurements of LDH release on the same samples used for the ELISA confirmed that macrophages were dying at 16 hours post-infection (Figure 6B). Threferore, although macrophages were dying, the IL-1β response to *C. auris* was minimal. In view of the low IL-1β response, we next compared the inflammatory potential of *C. auris* to that of *C. albicans*, side-by-side, using a virulent *C. albicans* isolate SC5314. *C. albicans* was used as a positive control here because it robustly activates the NLRP3 inflammasome (Gross et al., 2009; Hise et al., 2009; Joly et al., 2009; van de Veerdonk et al., 2011). As expected, *C. albicans* triggered high levels of IL-1ß secretion, in contrast to *C. auris* (Figure 6C). Therefore, in our experimental system, macrophages are clearly able to activate inflammasome-dependent responses when infected with *C. albicans*, but not with our *C. auris* isolate.

**Figure 6.**
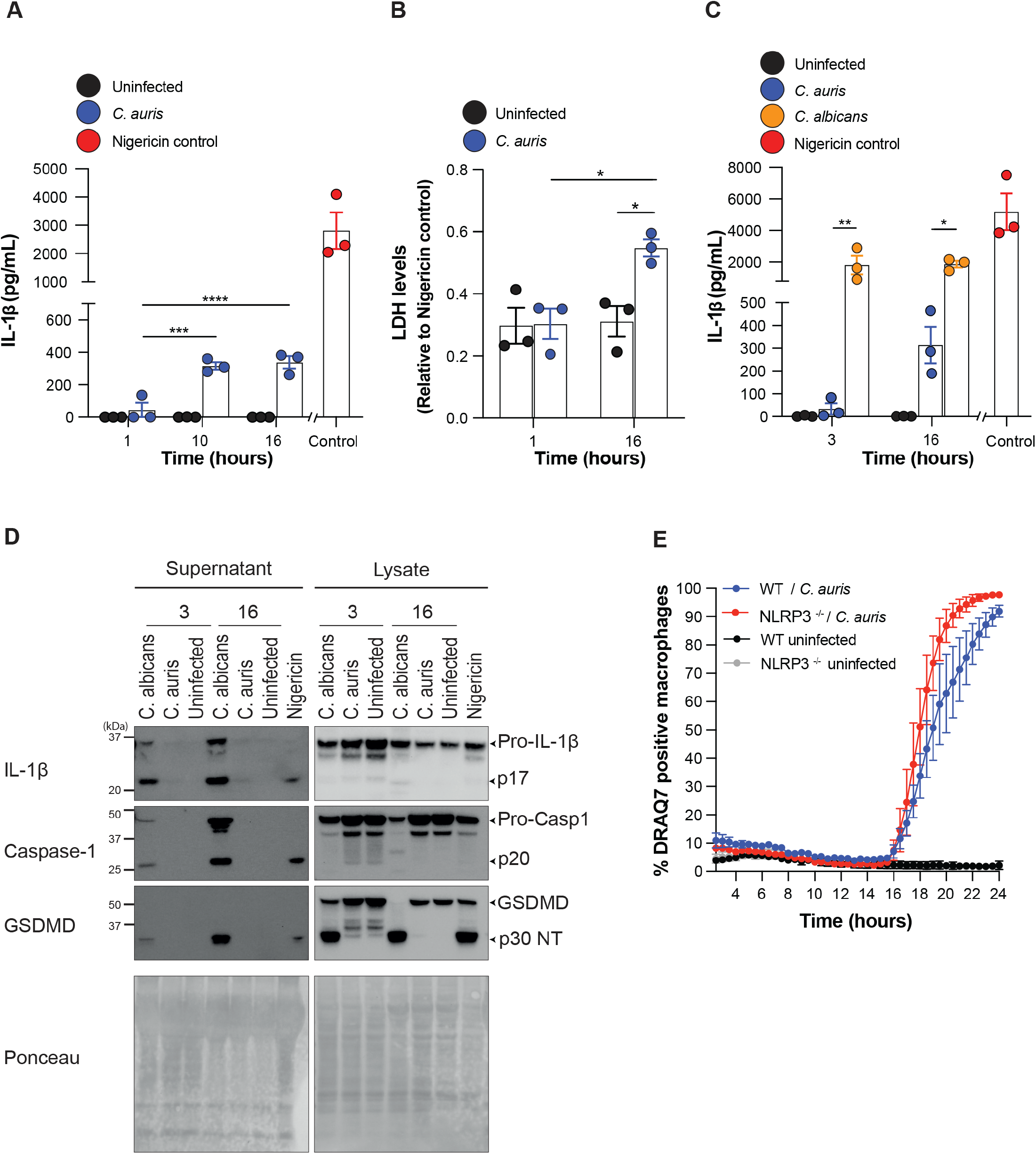
Macrophages fail to activate the NLRP3 inflammasome in response to *C. auris*. **A.** ELISA of IL-1β release in response to *C. auris* infection of BMDMs. Macrophages were primed with 50 ng/ml lipopolysaccharide (LPS) for 3 h, followed by challenge with *C. auris* (strain 470121) for 1, 10 and 16 h. The NLRP3 activator nigericin was used as a positive control and uninfected macrophages as a negative control. Shown are the mean values and SEM for 3 independent experiments. A 2-way ANOVA Bonferroni’s multiple comparison test was performed to compare the 10- and 16-hour infection condition to the 1-hour time point (*** p≤0.001, **** p≤0.0001). **B.** Lactate dehydrogenase (LDH) release for *C. auris-*infected and uninfected macrophages at 1 and 16 hours after challenge. The assay was performed on the supernatant samples used in panel A. All values are expressed relative to the nigericin control. Shown are the mean values and SEM from 3 independent experiments. A 2-way ANOVA Bonferroni’s multiple comparison test was performed to compare 16-hour infection samples to 1-hour time point (**** p≤0.0001). **C.** IL-1β ELISA was done as in panel A. Macrophages were primed with LPS as in A and challenged with *C. auris* (strain 470121), *C. albicans* (strain SC5314) or uninfected for 3 and 16 hours. Nigericin was used as a positive control. Shown are the mean values and SEM for 3 independent experiments. A 2-way ANOVA Bonferroni’s multiple comparison test was performed to compare the 10- and 16-hour infection condition to 1-hour time point (* p≤0.05, ** p≤0.01). **D.** Immunoblot analysis of supernatants and lysates from the same samples shown in panel C, at 3 or 16 hours after challenge. *C. albicans* and 10 *μ*M nigericin were used as positive controls and Ponceau staining acts as the loading control. The mature (active) IL-1β is the p17 fragment, for caspase it is the p20 fragment and p30 for Gasdermin D (GSDMD). A representative experiment is shown. Uncropped immuno blots for three independent experiments are shown in Figure S4. **E.** Macrophages were obtained from the bone marrow of wild type or *Nlrp3*^*-/-*^ mutant mice and infected with *C. auris* (strain 470121) with an MOI of 6. The appearance of DRAQ7-positive cells was quantified using live-cell imaging over 24 hours. Shown are the means and SEM from 3 independent experiments.

We next used biochemical approaches to directly test several further outputs of NLRP3 inflammasome activation. Western blot analyses of cellular lysates and supernatants largely confirmed the ELISA results, showing that *C. auris* failed to induce significant processing of IL-1β to its mature p17 form by the inflammasome-activated protease caspase-1 (Figure 6D, Figure S4). In comparison, *C. albicans* showed the expected induction of IL-1β processing (Figure 6D, Figure S4). We also examined two further markers of NLRP3 inflammasome activation: caspase-1 autoproteolysis (p20) and caspase-1-mediated cleavage of GSDMD to its N-terminal pore-forming fragment (p30) (He et al., 2015; Liu et al., 2016; Sborgi et al., 2016; Shi, J. et al., 2015). There was no evidence of caspase-1 or GSDMD cleavage upon *C. auris* infection (Figure 6D, Figure S4), further supporting the conclusion that the NLRP3 inflammasome is not responding. As expected, the control pathogen *C. albicans* induced both caspase-1 and GSDMD cleavage (Figure 6D, Figure S4). Finally, NLRP3 deficiency failed to reduce *C. auris*-induced macrophage cell death (Figure 6E). This result indicates that *C. auris* does not trigger NLRP3 inflammasome-dependent pyroptosis. Overall, our results suggest that *C. auris* avoids recognition by NLRP3, despite perturbing macrophage glycolysis and causing large-scale macrophage cell death.

## DISCUSSION

Our study establishes how *C. auris* evades macrophages and shows that the mechanisms are grounded in metabolic regulation. We show that *C. auris* escapes from immune containment and kills macrophages by causing host metabolic stress. *C. auris* escape and killing of macrophages are conserved mechanisms in isolates from four different pathogenic clades. Moreover, these mechanisms are recapitulated between mouse and human primary macrophages. Remarkably, our *C. auris* isolate is able to disrupt host glycolysis and kill macrophages while avoiding recognition by the NLRP3 inflammasome, which is in stark contrast with studies showing that disruption of macrophage glycolysis by *C. albicans* (Tucey et al., 2020), *Salmonella thyphimurium* (Sanman et al., 2016) and N-acetylglucosamine released from bacterial peptidoglycan (Wolf et al., 2016) activates the NLRP3 inflammasome and antimicrobial inflammation. Thus, *C. auris* is able to utilise metabolic regulation to eliminate macrophages without the associated costs that come from activating NLRP3 inflammasome-dependent antimicrobial responses. As such, we propose that this metabolic mechanism is a key immune evasion strategy that enables *C. auris* to survive and grow in infection niches. This proposition is supported by our data showing that deletion of the transcriptional activator *TYE7* inhibits glycolytic metabolism of *C. auris*, reduces its ability to kill macrophages and inhibits the establishment of infection foci in kidneys in the mouse bloodstream infection model. These findings also establish Tye7 as an important transcriptional regulator of *C. auris* metabolism and pathogenesis.

Our conclusion that *C. auris* egresses from macrophages and kills them contrasts with a recent study that showed no escape or macrophage damage by *C. auris* (Bruno et al., 2020). However, the authors only considered the first few hours after infection. By using our live-cell imaging platform that allows for dynamic assessment of host-pathogen interactions over prolonged infection times we demonstrate that escape does indeed occur, initiating at 8-10 hours after challenge. We further show that macrophage damage also occurs, initiating at 16-18 hours after challenge. Importantly, *C. auris* escape differs from that of the related fungal pathogen *C. albicans. C. albicans* egresses from macrophages by growing invasively in hyphal form and activating two pore forming proteins, fungal candidalysin and host gasdermin D (GSDMD) to damage macrophage membranes (Ding et al., 2021; Kasper et al., 2018; Lo et al., 1997; Olivier et al.). Our imaging shows that *C. auris* remains in yeast form as it escapes from macrophages and we also demonstrate biochemically that it does not trigger the processing of GSDMD to its pore-forming fragment (Figure 1 and 6). Furthermore, *C. auris* does not encode the fungal toxin candidalysin in its genome (Muñoz et al., 2018). Thus, unlike *C. albicans*, the act of *C. auris* escape is not associated with overt macrophage lysis.

Instead, our data supports a critical role for fungal metabolic capacity in *C. auris* immune escape. This conclusion is based on our results showing that the pyruvate kinase mutant *Δcdc19* is unable to escape from macrophages or kill them even when acetate is supplemented to support its growth (Figure 5). Metabolic capacity enables *C. auris* proliferation in macrophages, which could serve to increase pathogen loads to a sufficient degree to trigger non-lytic escape. The non-lytic escape from macrophages might also provide clues as to why there is no NLRP3 inflammasome activation in response to *C. auris*. NLRP3 activation usually requires two signals. Based on work with *C. albicans*, Signal 1 is provided by recognition of fungal cell wall components by macrophage receptors, while Signal 2 is provided by rupture of the phagosomal membrane by hyphal invasive growth (Gross et al., 2009; Hise et al., 2009; Westman et al., 2020). Hiding of β-glucan by *C. auris* mannan has been proposed to prevent immune recognition (Wang et al., 2022). This could reduce Signal 1. Of note however, in our experiments that assess inflammasome outputs (Figure 6) macrophages are pre-treated with LPS. LPS provides Signal 1, yet we show that inflammasome activation still does not occur. Therefore, we believe that lack of Signal 1 is not the explanation. Our data that *C. auris* remains in yeast form and escapes by a non-lytic mechanism suggests that the phagosomal membrane remains intact during escape. This means that there is no Signal 2, which could prevent the priming of macrophages for activation of NLRP3 even upon metabolic dysfunction and lytic cell death that we show occur later in *C. auris* infection. In this way, the metabolic adaptations that enable *C. auris* to grow in macrophages and trigger non-lytic escape could also contribute to its hiding from immune recognition.

On the host side, we present evidence that *C. auris*-infected macrophages undergo immunometabolic shifts to increase their glycolytic metabolism. Increased macrophage glycolysis is a conserved response to bacterial and fungal pathogens, as well as microbial ligands (Domínguez-Andrés et al., 2017; Gonçalves et al., 2020; Shi, L. et al., 2015; Tannahill et al., 2013; Tucey et al., 2018). Surprisingly however, *C. auris-*infected PBMCs did not display transcriptional signatures of increased glycolysis and it was therefore suggested that this metabolic response of immune phagocytes might be different with *C. auris* (Bruno et al., 2020). Our data challenges this conclusion and we show that glycolysis is critically important for *C. auris*-infected macrophages, as macrophages were rescued from *C. auris*-induced cell death by glucose and pyruvate (the terminal metabolite of glycolysis), but not by acetate, glutamine or arginine that feed into the TCA cycle at different points (Figures 3 and 4). Differences in experimental conditions, including distinct *C. auris* isolates and immune cell types (PMBC versus BMDMs) could explain these contrasting conclusions. We note that we recapitulated immune cell killing of *C. auris* in both human and mouse macrophages and show that macrophage killing depends on their reliance on glucose, which is a consequence of the immunometabolic shift. Therefore, our data suggest that both human and mouse macrophages undergo immunometabolic reprogramming in response to *C. auris*, which renders them susceptible to glucose competition by *C. auris* that causes immune cell death.

We found a key difference between *C. auris* and previously studied pathogens with respect to the roles of macrophage glycolysis in antimicrobial responses. Increased glycolysis contributes to antifungal immune responses and cytokine production by macrophages and monocytes in response to *C. albicans* and *Aspergillus fumigatus* (Domínguez-Andrés et al., 2017; Gonçalves et al., 2020) and also in response to several bacterial pathogens (Gleeson et al., 2016; Huang et al., 2018; Tannahill et al., 2013). However, we show that these functions are not recapitulated with *C. auris*. We detected very little IL-1β secretion by *C. auris-*infected macrophages and minimal signs of processing of this cytokine to its active form by the NLRP3 inflammasome. Our conclusion is supported by a study published while our manuscript was in preparation, and which also showed low IL-1β secretion by *C. auris*-infected mouse macrophages (BMDMs)(Wang et al., 2022). In contrast, a recent report with human PBMCs concluded that *C. auris*-induced cytokine responses were more potent relative to *C. albicans* (Bruno et al., 2020). Differences in cytokine secretion could reflect the use of distinct *C. auris* isolates and immune cell types (Wang et al., 2022). Wang et al attributed low cytokine responses to reduced recognition and phagocytosis of *C. auris* by macrophages, but we see effective phagocytosis. Our data shows that a key mechanism causing low IL-1β responses is that *C. auris* is not competent to activate the NLRP3 inflammasome. Specifically, we could not detect any of the known outputs of NLRP3 inflammasome activation, as there was essentially no processing of IL-1β, caspase-1 or GSDMD and no activation of pyroptosis in *C. auris*-infected macrophages. Taken together, our findings argue that, although macrophages upregulate glycolysis in response to *C. auris*, this does not serve the usual roles of promoting IL-1β cytokine production and supressing microbial growth.

In conclusion, our findings establish important mechanisms of macrophage evasion by *C. auris* involving metabolic adaptations in the pathogen that promote egress from the immune phagocytes and creation of a nutrient-poor infection niche in which macrophages cannot survive. Unexpectedly, macrophage cell death and disruption of host glycolysis by our *C. auris* isolate did not activate the NLRP3 inflammasome. Thereby, macrophage IL-1β responses and their ability to recruit immune cells such as neutrophils into infection sites, remain low. Given the contrasting conclusions on IL-1β responses to *C. auris* from various studies to date, future studies should address inflammasome responses to the broadest possible range of *C. auris* clinical isolates. Neutrophils also need glucose for antifungal responses, as has been demonstrated comprehensively for *C. albicans* in a recent study (Li et al., 2022). *C. auris* is poorly killed by neutrophils (Johnson et al., 2018) and it will be interesting to determine if metabolic mechanisms and glucose deprivation play a role in its neutrophil evasion. Collectively, our study suggests that manipulation of metabolism could be an effective strategy to improve the ability of innate immune phagocytes to control *C. auris* and curb yeast growth in host tissues to limit infection. Given that *C. auris* can be resistant to all current classes of antifungal drugs, it is important to find alternative treatment strategies. Deeper understanding of metabolism in *C. auris* infections contributes to this goal.

## MATERIALS AND METHODS

### Experimental model

#### *Candida* strains and media

The strains used in this study are listed in Table S1. The *C. auris* isolate 470121 was generously gifted by Sarah E. Kidd from the National Mycology reference centre (Adelaide, Australia) and is a clinical isolate obtain from Dr. Sharon Chen. The other *C. auris* clinical strains used in this study were obtained from the Centre for Disease Control (CDC, Atlanta, USA). The *C. albicans* clinical isolate was SC5314. All strains were maintained on Yeast Extract – Peptone – Dextrose (YPD) media (1% yeast extract, 2% peptone, 2% glucose, 80 μg/ml uridine, with addition of 2% agar). Where overnight liquid cultures were required such as for macrophage infections or growth curve assays, a single colony of *Candida* from a freshly streaked YPD plate was inoculated into liquid YPD medium and grown for 18 hours at 30 °C. For liquid growth assays, overnight cultures of *C. auris* were diluted to OD_600nm_ of 0.1 in prewarmed bone marrow derived macrophage (BMDM) medium containing RPMI 1640 supplemented with 12.5 mM HEPES, 10% FBS, 20% L-cell conditioned medium (containing macrophage colony-stimulating factor) and 100 U/ ml of penicillin-streptomycin (Sigma) (complete RPMI media). For growth assays investigating the effects of different types and concentrations of carbon sources, BMDM medium was made up with RPMI 1640 medium lacking glucose (Thermo Fisher 11879020) and supplemented with glucose (10, 20 and 40 mM), galactose (40 mM) or mannose (40 mM). For yeast plate growth tests on various nutrient sources, 10-fold dilutions of yeast cultures were spotted on minimal YNB media was supplemented with 2% of either glucose, fructose, mannose, acetate or glycerol. For some growth assays, 0.5 *μ*g/ml antimycin was supplemented to the plates.

#### Strain construction

The *Δtye7* and *Δgal4* mutant strains were generated from clade I strains 470121 and B8441, and the *Δcdc19* mutant strain was generated in the B8441 strain. Mutants were produced by deletion of the open reading frame using the SAT1 flipper cassette (Reuss et al., 2004), conferring resistance to nourseothricin. Primers used to generate the knockout cassettes that enabled the generation of these mutants can be found in Table S2. Deletion mutants were selected on YPD media supplemented with 400 *μ*g/ml nourseothricin. The mutants were generated by standard methods based on PCR and homologous recombination, however as *C. auris* has a lower rate of homologous recombination compared to other *Candida* species (Santana and O’Meara, 2021), W7 Hydrochloride (5*μ*g/ml) was added to the growth culture as detailed in (Arras and Fraser, 2016), to reduce non-homologous end joining. All strains were confirmed by diagnostic PCR and primers used for this can be found in Table S2. Both parent strains 470121 and B8441 are prototrophic and nourseothricin sensitive.

#### Isolation of murine bone marrow derived macrophages (BMDMs) and human monocyte-derived macrophages (hMDMs)

Mice experiments for BMDM isolations were approved by the Monash University Animal ethics committee (approval numbers ERM14292 and ERM25488). Human blood isolation experiments were approved by the Monash University Human Ethics Committee (project 21685).

For BMDM extractions, tibia and femur were harvested from six to eight weeks old C57BL/6 mice obtained from the Monash Animal Research Platform. For experiments using *Nlrp3* ^*-/-*^ BMDMs, tibia and femur from *Nlrp3* ^*-/-*^ mice were obtained from the Walter and Elisa Hall Institute. Bone marrow containing monocytes was flushed from these bones using BMDM media (RPMI 1640 supplemented with 12.5 mM HEPES, 10% FBS, 20% L-cell conditioned medium and 100U/ ml of penicillin-streptomycin) and allowed to differentiate into BMDMs for 5-7 days as previously described(Tucey et al., 2016).

For isolation of hMDMs, blood of healthy volunteers was used to isolate PBMCs by Histopaque-1077 (Sigma-Aldrich) density gradient centrifugation. Using the Pan Monocyte Cell Isolation (Miltenyi Biotech) magnetic labelling system, CD14 and CD16-expressing monocytes were purified from the PBMCs. This enriched monocyte fraction was differentiated to macrophages (hMDMs) in a 25 cm^2^ culture flasks in RPMI 1640 media containing 15 mM HEPES, 10% fetal bovine serum (Serana, Fisher Biotech), 100 U/L penicillin-streptomycin (Sigma-Aldrich) and 50 ng/mL macrophage colony-stimulating factor (M-CSF, R&D Systems) and incubated at 37°C in 5% CO_2_ for 7 days.

### Method details

#### Live Cell Imaging

For seeding macrophages in live cell imaging experiments, cells were gently scraped from petri dishes using a cell scraper (BD Falcon) and seeded in tissue culture-treated plates at a density of 1 × 10^5^ cells/well in a 96-well tray and incubated overnight at 37°C + 5% CO_2_. Live cell imaging procedure was followed as described in(Tucey et al., 2016). Briefly, macrophages were stained with 1 mM CellTracker Green CFMDA dye (ThermoFischer C7025) for 35 minutes in serum-free RPMI 1640 supplemented with 12.5 mM HEPES, 20% L-cell conditioned medium and 100 U/ ml of penicillin-streptomycin (Sigma). Macrophages were then infected with *C. auris* at a multiplicity of 3:1 or 6:1 in BMDM medium and co-incubated for 1 hour, followed by removal of non-phagocytosed yeast by washing with PBS. In some experiments, BMDM media was supplemented with arginine, glutamate, acetate, pyruvate, sorbitol or glucose at the concentrations indicated in the figures. For experiments investigating the effects of glucose, concentrations BMDM media was prepared with RPMI 1640 medium lacking glucose (Thermo Fisher 11879020) and 10 % FBS. This media was used as un-supplemented (“no glucose”) or supplemented with 10, 20 or 40 mM glucose. Where indicated, 125 ng/ml caspofungin and added post co-infection. For experiments involving metformin, macrophages were pre-treated with 5 mM metformin for 24 hours and then switched to fresh media with or without metformin prior to infection with *C. auris*. Uninfected macrophage controls were treated similarly. All media conditions also contained 0.6 mM DRAQ7 (Abcam) for tracking macrophage membrane permeabilisation. Imaging was performed on a Leica AF6000 LX epifluorescence microscope (20x objective). The imaging data was analysed and quantified using MetaMorph (Molecular Devices) as previously described (Tucey et al., 2016) and plotted using Prism 9.0 (GraphPad Software, San Diego, California USA) software.

For experiments with hMDMs, cells were detached in PBS with 2 mM EDTA and 2 % FBS by incubating for 10 minutes at 37 °C in 5% CO_2_. This was followed by repeated rinses using a 10 mL syringe and 18G needle. The detached hMDMs were then seeded at a density of 5 × 10^4^ cells/well of a 96-well plate, in RPMI 1640 media supplemented with 15 mM HEPES, 10 % fetal bovine serum (Serana, Fisher Biotech), 100 U/L penicillin-streptomycin and incubated at 37 °C and 5% CO_2_ overnight before infection. hMDMs were stained with 1 mM CellTracker Green CFMDA dye (ThermoFischer C7025) for 30 minutes prior to infection as above for BMDMs. After staining, hMDMs were primed in RPMI 1640 supplemented with 15 mM HEPES, 10% and 50 ng/ml lipopolysaccharide (LPS, Sigma) and incubated at 37 °C and 5% CO_2_ for 2 hours for priming. After priming hMDMs were challenged with *C. auris* (strain 470121) at MOI of 6:1 (*Candida*: macrophage) for 1 hour and the remaining procedures for live cell imaging were as described above for BMDM experiments.

#### Collection of supernatants and lysates for IL-1β ELISA, Lactate dehydrogenase (LDH) and Western blot assays

The seeding of macrophages was performed as described in Live cell imaging methods. All assays were carried out in 24-well microplates and macrophages were seeded at 5 × 10^5^ cells/well in BMDM media. Macrophages were primed with 50 ng/ml Lipopolysaccharide (LPS) for 3 hours in fresh media, followed by either challenge with *C. auris* (strain 470121, MOI 6), *C. albicans* (strain SC5314 MOI 3) or uninfected for 1 hour at 37 *°*C and 5% CO_2_. After this, cells were washed 3x in PBS to remove non-phagocytosed fungal cells, replenished with 250 *μ*l fresh BMDM media and incubated for 3, 10 and 16 hours. The 1-hour time point was sampled directly after infection incubation. Treatment with 10 *μ*M nigericin for 3 hours was used as a positive control. For each time point, supernatants were collected for ELISA (100 *μ*l), western blot (75 *μ*l) and LDH assay (75 *μ*l). ELISA and LDH assay supernatants were stored at -80°C. Western blot supernatants were mixed with 25 *μ*L 4x sample buffer (RSB; ThermoFisher) + 5% β-mercaptoethanol in microfuge tubes, boiled at 95 *°*C for 10 min and frozen at -20 *°*C. After collection of the supernatants, 150 *μ*L of 1x RSB buffer + 2.5 % β- mercaptoethanol was added to each well and a 1 *μ*l pipette tip was used to scrape cells for lysate faction. This was transferred to 1 microfuge tubes, boiled at 95 *°*C for 10 min and frozen at -20 *°*C.

#### IL-1β ELISA

The IL-1β levels in supernatant samples were measured using the Mouse IL-1β/IL-1F2 ELISA kit (R&D Systems, DY401), according to the manufacturer’s instructions. Optical density measurements reflective of IL-1β levels were measured at 450 nm (with a wavelength correction of 540 nm) using a Tecan infinite M200 plate reader. Using the reference of IL-1β standards provided in the kit, an interpolated standard curve was generated in Graphpad Prism 9.0 (GraphPad Software, San Diego, California USA) to determine IL-1β levels for each sample.

#### LDH assay

The LDH levels in supernatant samples were measured using the CytoTox 96 Non-23 Radioactive Cytotoxicity Assay (Promega, G1780) according to the manufacturer’s instructions. Optical density measurements reflective of LDH levels were measured at 492 nm using a Tecan Spark 10M plate reader. LDH levels for each sample were quantified as fold change relative to the 10 *μ*M nigericin positive control.

#### Western blot

Prior to loading, cell lysates and/or supernatants were boiled for 5-10 minutes at 95°C, followed by loading onto 4-12% Bis-Tris NuPage protein gels (Invitrogen), SDS-PAGE and transfer onto nitrocellulose membrane (Millipore). Membranes were stained with Ponceau. Blocking was with 5% (v/v) skim milk in PBS-Tween (0.1 % (v/v)) for 1 hour, followed by probing with primary antibodies: anti-IL-1β (R&D, #AF-401-NA), anti-caspase-1 (Adipogen, #AG-20B-0042-C100) and anti-GSDMD (abcam, ab209845). Incubations with primary antibodies were overnight in a rotating wheel at 4°C, followed by washing in PBS-T and adding HRP-conjugated secondary antibodies (1 hour at room temperature). Membranes were developed using Immobilin Forte Western HRP (Merck) and imaged using ChemiDoc (BioRad). Analyseis was performed using Image Lab and Adobe Illustrator.

#### Lactate assay

The seeding of macrophages was as described in Live cell imaging methods and assays were carried out in 96-well microplates with macrophages seeded at 1 × 10^5^ cells/well in BMDM media. Macrophages were either challenge with *C. auris* (strain 470121, MOI 6) or uninfected for 1 hour at 37 *°*C and 5% CO_2_. Following the 1-hour infection, cells were washed thrice in PBS, replenished with 150 *μ*l fresh BMDM media and incubated for 10 and 24 hours. The 1-hour time point was sampled directly after infection. At each time point, the supernatant was removed, plates washed 4 times in cold PBS and cells harvested in 4x Lactate Assay buffer (about 200 *μ*l). The preparation of samples and lactate level measurement procedure were followed as specified in the Lactate Assay Kit (Colorimetric) (ABCAM - ab65331) according to the manufacturer’s instructions. Optical density was measured at 450 nm using a Tecan Spark 10M plate reader. Using the reference of lactate standards provided in the kit, an interpolated standard curve was generated in Graphpad Prism 9.0 (GraphPad Software, San Diego, California USA) to determine lactate levels for each sample.

#### Quantification of glucose

To quantify glucose in *C. auris*-macrophage infection assays, supernatant was collected at 1, 4, 8 10 and 24 hours from macrophages (96-well microplates seeded at 1 × 10^5^ cells/well) either challenge with *C. auris* (strain 470121, MOI 6) or uninfected. Glucose levels were measured using the Amplex Red glucose/glucose Oxidase Assay Kit (Thermo Fisher A22189) according to the manufacturer’s instructions. Optical density measurements were carried out at 450 nm using a Tecan Spark 10M plate reader.

#### Colony forming unit (CFU) measurements

For counting CFUs, macrophages were seeded as described in Live cell imaging methods. All assays were carried out in 24-well microplates and macrophages were seeded at 2.5 × 10^5^ cells/well in BMDM media. *C. auris* cultures were grown over night for 18 hours in YPD medium at 30 *°*C. The next day macrophages were challenged with *C. auris* (MOI 3 for 1 hour). Cells were then washed 3x in PBS, replenished with 250 *μ*l fresh BMDM media and incubated for 2, 4, 8 and 10 hours. At each time point supernatant was removed and saved, and macrophages were lysed with ice-cold distilled water to obtain intracellular *C. auris*. Appropriate dilutions of supernatant (extracellular *C. auris*) and lysate (intracellular *C. auris*) were plated on YPD plates and incubated for 2 days at 30 *°*C to count yeast colonied.

#### Growth curves

For growth curve assays WT, *Δtye7, Δgal4* and *Δcdc19* strains were cultured as described in the *Candida* strains and media method. Assays were run in either BMDM media or BMDM media with 10 mM glucose or acetate. A matched media only controls were also run alongside the *C. auris* samples. Assay was conducted in 200 *μ*l volumes of a 96-well plate. Growth was determined by optical density measurements at 600 nm using a Tecan Spark 10M plate reader over a 24-hour period. Doubling times were calculated using the Growthcurver package in Rstudio (https://cran.rproject.org/web/packages/growthcurver/vignettes/Growthcurvervignette.html#output-metrics)(Sprouffske and Wagner, 2016).

#### Phagocytosis determinations

Rates of *C. auris* phagocytosis by macrophages were calculated using images captured from experiments shown in Figure 1A. Briefly, a random sampling of macrophages in a fixed area was used in counts from images for n=3 independent replicates. More than 100 macrophages were counted for each replicated and included both *C. auris*-infected and uninfected cells. The number of *C. auris* in each macrophage was counted. The phagocytic ratio was calculated by dividing the total number of counted *C. auris* by the total number of macrophages. The data from the 3 independent experiments is presented in Dataset S1.

#### Quantitative PCR analysis (qPCR)

Primers used for qPCR are detailed in Table S2. For RNA isolation in *C. auris*/macrophage infections BMDMs were seeded at a density of 1 × 10^6^ cells/well in a 6-well tray, and then infected with *C. auris* at a MOI of 6 in BMDM medium. After 1 hour of co-incubation and PBS washes, media was replaced for each well with fresh BMDM medium. Macrophage samples without *C. auris* infection (control samples) were treated in the same way. Samples were incubated for 1, 4, 8 and 10 hours. At the given time points, samples were harvested using Trizol reagent to lyse the cells. The lysate was centrifuged to separate the *Candida* pellet from the macrophage RNA supernatant in the infected samples. RNA from all samples was isolated using the Trizol method. Reverse transcription was carried out on 1 μg of DNase I (Ambion)-treated total RNA using Superscript III cDNA kit (Invitrogen). Quantitative PCR was performed on a LightCycler 480 (Roche) using the FastStart Universal SYBR Green Master Rox (Roche) master mix. Data analysis was carried out using the LinReg software(Ruijter et al., 2009) and macrophage glycolytic gene expression was normalised to 18S rRNA levels. Two technical repeats were averaged for one biological repeat.

For qPCR experiments assessing *C. auris* glycolysis gene expression, yeast cultures were grown *in vitro* at 37 *°*C. Overnight cultures = were diluted to OD_600nm_ 0.1 in BMDM media and grown at 37 *°*C for 7 hours. Total RNA was isolated using the hot phenol method and DNAse treatment, reverse transcription, qPCR method and data analysis were carried out as described above. *RDN25* transcript levels were used for normalisation. Two technical repeats were averaged to obtain the biological repeat.

#### Animal infections

Six-week old female BALB/c mice weighing 18 to 22 g were purchased from Envigo, Rehovot, Israel. Animal experiments were reviewed and authorized by the Tel Aviv Sourasky Medical Center Institutional Animal Care and Use committee (permit number TLVMC - IL - 2112 - 124 – 3). Mice were housed in filter topped cages and fed autoclaved food and water. *C. auris* wild type and *Δtye7* strains were grown in liquid yeast-extract agar glucose medium (YAG; yeast extract at 5 g/liter; glucose at 10 g/liter; agar at 15 g/liter; 1 M MgCl_2_ at 10 ml/liter; trace elements at 1 ml/liter) in a shaking incubator at 30 °C, washed twice in sterile PBS, and resuspended in PBS at a concentration of 10^8^ cells/ml (inoculum suspension). For each tested *C. auris* strain, mice were infected in groups of 7 with 10^7^ yeast cells in 100 µl PBS by lateral tail vein injection. Mice were euthanised using CO_2_ inhalation 48 and 120 hours after infection. Kidneys were excised aseptically, weighed and homogenized in 1 ml sterile PBS. Homogenates were serially diluted in PBS, plated on YAG and incubated at 30°C for 48 hours. CFU counts were used to calculate the tissue fungal burden (CFU/g).

#### Statistical analysis

Statistical analyses were conducted in Graphpad Prism 9.0 (GraphPad Software, San Diego, California USA) and is detailed in each of the figure legends. 2-way ANOVA Bonferroni’s multiple comparison tests were conducted on CFU counts, LDH assays, Lactate level assay, IL-1β assay, ELISA experiments and qPCR experiments. Kruskal Wallis non-parametric test and the Dunn’s multi comparison test was conducted on the fungal qPCR assay and assays determining fungal burden in animal infection. For all experiments a P value of less than 0.05 was considered to be statistically significant.

## SUPPLEMENTARY FIGURE LEGENDS

**Supplementary Figure 1. *C. auris* replicates inside macrophages, followed by egress**

**A.** Images of *C. auris*-infected macrophages from the live cell imaging. These are the entire field of view for the images shown in Figure 1B. The parts of the image shown in Figure 1B are indicated by a perforated box. The yellow arrowhead shows escaped *C. auris*.

**B.** Quantitation of intracellular (left panel) and extracellular (right panel) *C. auris* colony forming units (CFUs) during BMDM challenge in the presence of glucose, galactose, mannose (40mM) and no carbon source supplementation. BMDMs were challenged with *C. auris* strain 470121 at MOI of 3 and both supernatant and macrophage lysate plated at 2, 4, 8 and 10 hours after challenge. CFUs were counted after 2 days of incubation at 30 °C. The bar graph shows the mean values and standard error of the mean (SEM) from n=3 independent experiments. Statistical significance was determined using a 2-way ANOVA Bonferroni’s multiple comparison test was used ((*p≤0.05, ** p≤0.01, ***p≤0.001, **** p≤0.0001). Only statistically significant comparisons are indicated in the figure.

**C.** *C. auris* growth *in vitro* in the presence of various carbon sources. Overnight cultures of *C. auris* (strain 470121) were diluted to an OD_600_ of 0.1 in media with no carbon source or 40 mM of either glucose, galactose or mannose and optical density measured at 1hour intervals over 24 hours. Shown here are the mean values and SEM for n=3 independent experiments.

**Supplementary Figure 2. Growth phenotypes of *C. auris* clinical isolates**

**A.** *In vitro* growth of *C. auris* isolates in BMDM media containing glucose. Overnight yeast cultures were diluted to an OD_600_ of 0.1 and optical density measured at 1-hour intervals over 24 hours. Each trend line represents an isolate from a discrete Clade: Clade I: B11098, B8841 and 470121, Clade II: B11220, Clade III: B11221 and Clade IV: B11244. Shown here are the mean values and SEM for 3 independent experiments.

**B.** *In vitro C. auris* growth in the presence of varying concentrations of glucose (10 mM, 20 mM and 40 mM). Overnight cultures of *C. auris* (strain 470121) were diluted to an OD_600_ of 0.1 in media with the indicated concentrations of glucose. Carbon source free media was also used (no added glucose). OD_600_ was measured at 1-hour intervals over 24 hours. Shown are the mean values and SEM for 3 independent experiments.

**Supplementary Figure 3. Growth phenotypes of the *C. auris* glycolytic mutants**

**A.** Liquid growth assays of the indicated strains (wild type and pyruvate kinase mutant *Δcdc19*) in medium containing glucose or acetate over 24 hours in BMDM media. Shown are the means and SEM of n=3 experiments.

**B.** Liquid growth assays of wild type, *Δtye7* and *Δgal4* in glucose containing BMDM media. The strain background is indicated above the graph. Shown are the means and SEM of n=3 experiments.

**C.** Doubling time calculations for wild type, *Δtye7* and *Δgal4* from liquid growth assay in C. Shown are the means and SEM for n=3 independent experiments. ****p≤0.0001 (2-way ANOVA Bonferroni’s multiple comparison test). Only statistically significant comparisons are indicated in the figure.

**Supplementary Figure 4. Individual Western blot experiments**

Independent repeats of Western blots shown in Figure 7D. Ponceau staining was used a loading control. Experiment 3 is the repeat shown in Figure 5D.

**Supplementary Movie 1. *C. auris* escapes macrophages and undergoes rapid extracellular replication**

Live cell imaging of *C. auris* yeast cells escaping from macrophages between 8 to 10 hours after challenge. The video is of the same field of view presented in Figure 1B. DRAQ7 staining (red dot signals in the image) indicates macrophages that have their cell membrane permeabilised, as an indication of cell death. MOI is 3:1 (*C. auris*: macrophage) and scale bars represent 100 *μ*m. Image brightness and contrast were adjusted as described in Materials and Methods.

**Supplementary Movie 2. Example of *C. auris* escaping from macrophages**

Live cell time-lapse of *C. auris* yeast cells escaping from a macrophage between 7 to 9 hours after challenge. Images were captured in 10 min intervals. DRAQ7 staining indicates macrophages that have died (i.e., cell membrane has been disrupted) and is signified by a red dot signal in the image. MOI is 3:1 (*C. auris*: macrophage) and scale bars represent 20*μ*m. Image brightness and contrast were adjusted as described in Materials and methods.

**Supplementary Table 1. *C. auris* strains used in this study**

**Supplementary Table 2. Primers used in this study**

**Supplementary Dataset 1. Data used to construct figures**

## ACKNOWLEDGMENTS

We thank Sarah Kidd (Australian Mycology Reference Centre) and Sharon Chen for the *C. auris* strain 470121 and Anastasia Litvintseva from the CDC for their panel of *C. auris* strains. We further thank James Vince and Seth Masters (Walter and Eliza Hall Institute) from providing *Nlrp3*^*-/-*^ macrophages and the Monash MicroImaging Facility (MMI) for expert support with microscopy.

This work was supported by grants from the Australian National Health and Medical Research Council (NHMRC) (APP1158678 to A.T., APP2002520 to A. T. and R.B-A. and APP1181089 to K.E.L.). R. B-A is further supported by Israeli Science Foundation grant 442/18 and Ministry of Science and Technology grant 88555. C.S. is supported by a fellowship from the Monash-Warwick Alliance AMR Training Program. A.T. and K.E.L. are Future Fellows of the Australian Research Council (FT190100733 to A.T. and FT19010266 to K.E.L.).

## AUTHOR CONTRIBUTIONS

H.W., C.S., R.B-A., and A.T. designed the study. H.W., C.S., T.D., I.T., T.L.L., D.S., N.M., F.A.B.O., and M.S. performed the experiments. H.W., C.S., D.S., and N.M., analysed the data. R.B-A., K.E.L., and A.T. provided advice on experimental approach and data interpretation. A.T., K.E.L. and R.B-A obtained funding. A.T. wrote the manuscript with contributions from H.W. and editorial contributions from all authors.

## Notes

### Competing Interest Statement

The authors have declared no competing interest.

