## Supplementary Figures and Tables for "*Candida auris* evades innate immunity by using metabolic strategies to escape and kill macrophages while avoiding antimicrobial inflammation"

#### Figure S1

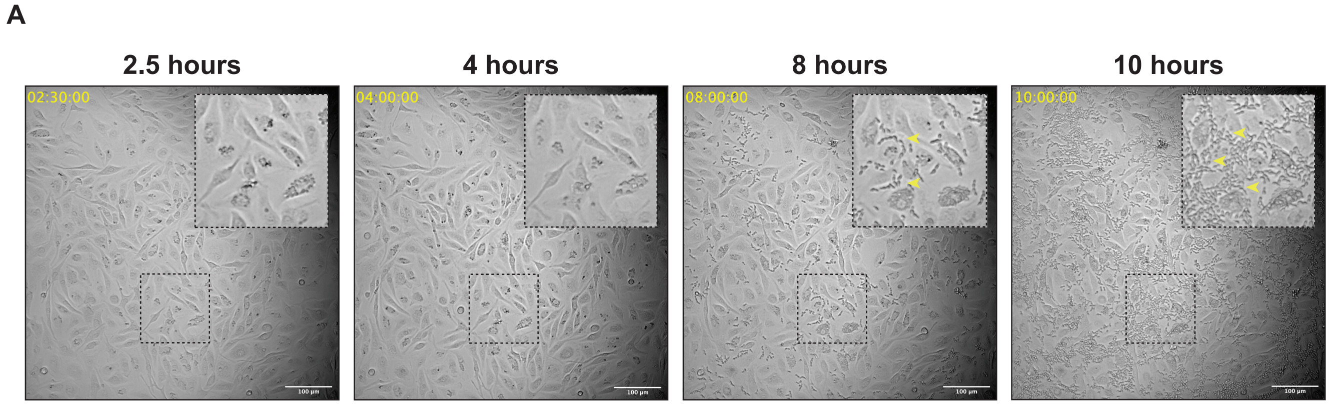

A
2.5 hours
4 hours
8 hours
10 hours
02:30:00
04:00:00
08:00:00
10:00:00


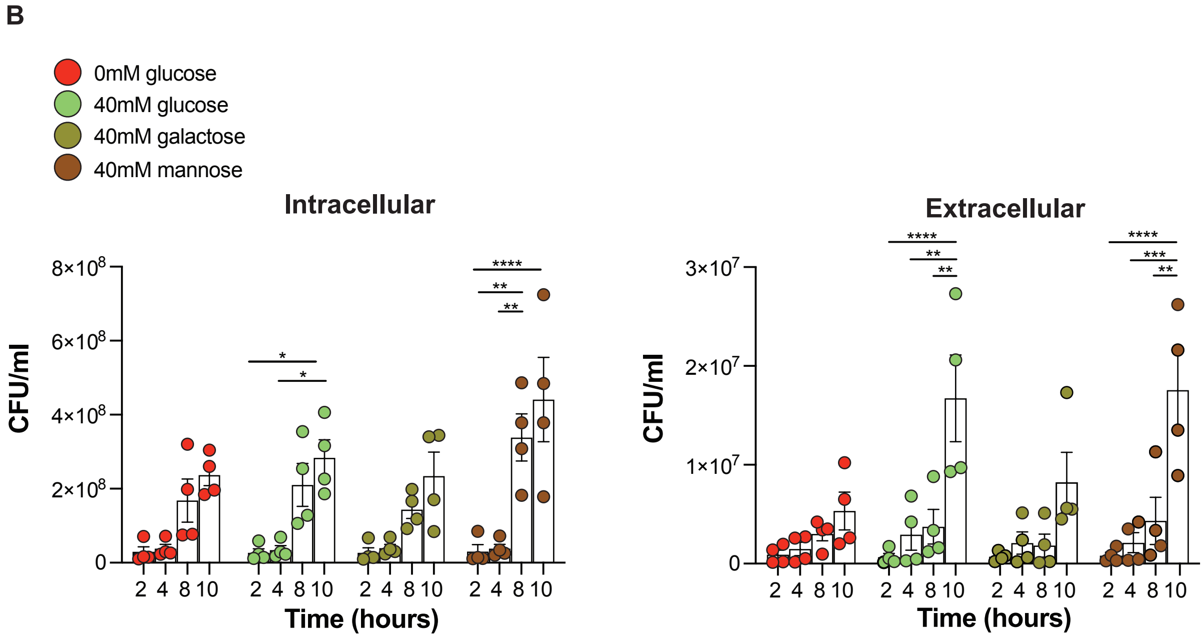

B
0mM glucose
40mM glucose
40mM galactose
40mM mannose
Intracellular
Extracellular
CFU/ml
Time (hours)


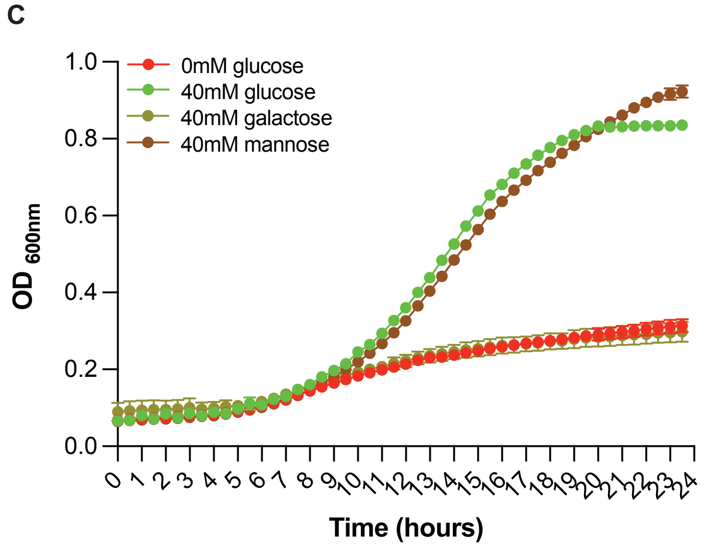

C
0mM glucose
40mM glucose
40mM galactose
40mM mannose
OD 600nm
Time (hours)

#### Figure S2

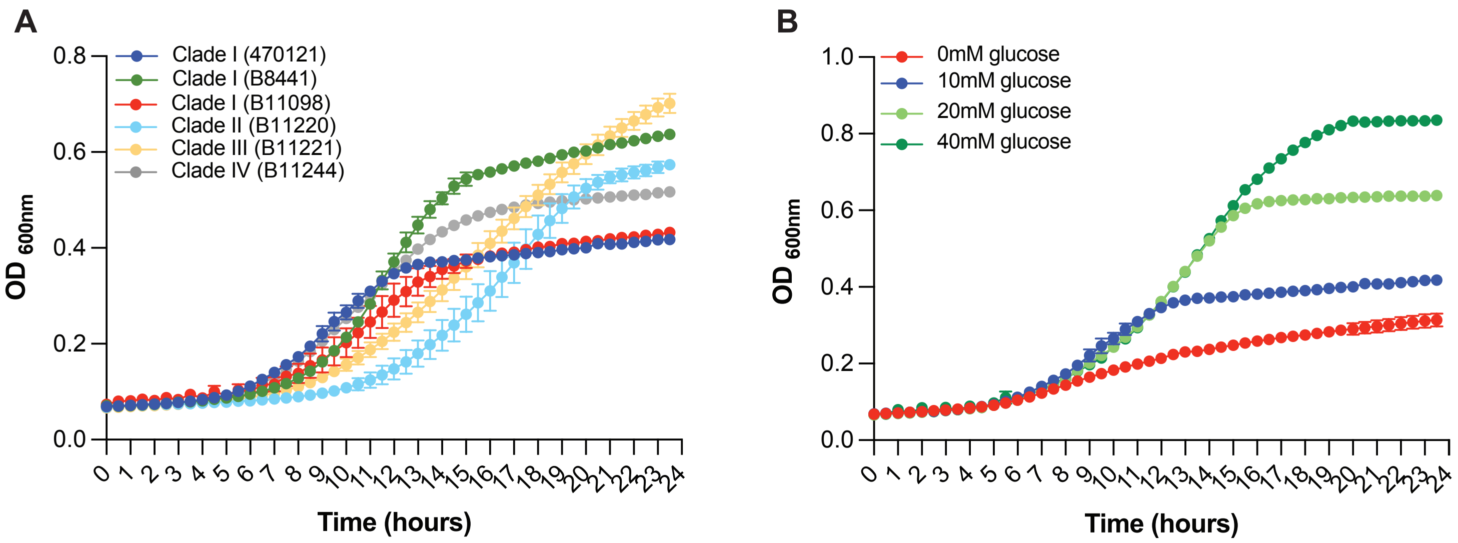

A
Clade I (470121)
Clade I (B8441)
Clade I (B11098)
Clade II (B11220)
Clade III (B11221)
Clade IV (B11244)
OD 600nm
Time (hours)
B
0mM glucose
10mM glucose
20mM glucose
40mM glucose
OD 600nm
Time (hours)

### Figure S3

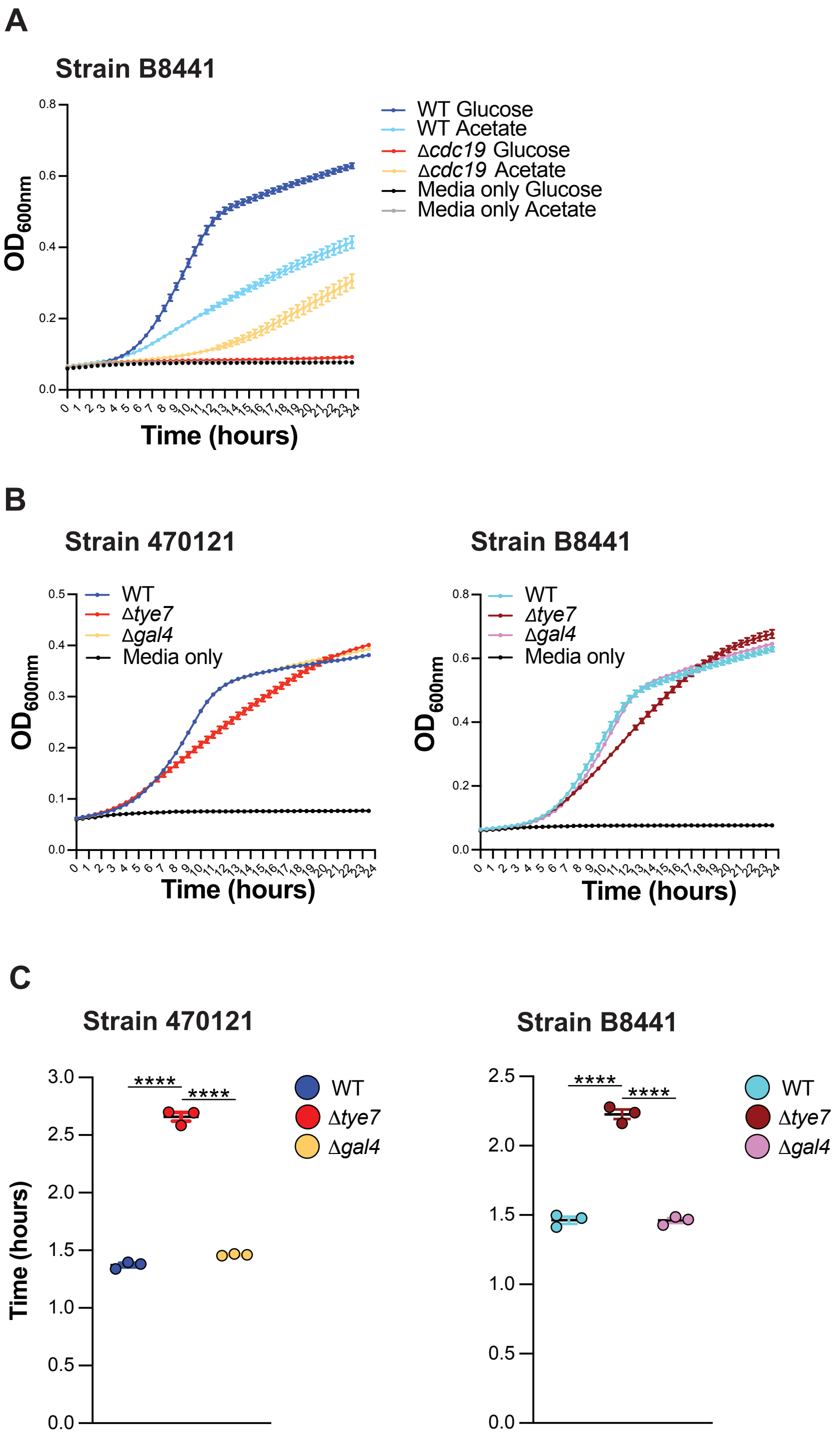

A
Strain B8441
WT Glucose
WT Acetate
Δcdc19 Glucose
Δcdc19 Acetate
Media only Glucose
Media only Acetate
OD600nm
Time (hours)
B
Strain 470121
WT
Δtye7
Δgal4
Media only
OD600nm
Time (hours)
Strain B8441
WT
Δtye7
Δgal4
Media only
OD600nm
Time (hours)
C
Strain 470121
Time (hours)
****
****
WT
Δtye7
Δgal4
Strain B8441
****
****
WT
Δtye7
Δgal4

#### Figure S4

### A

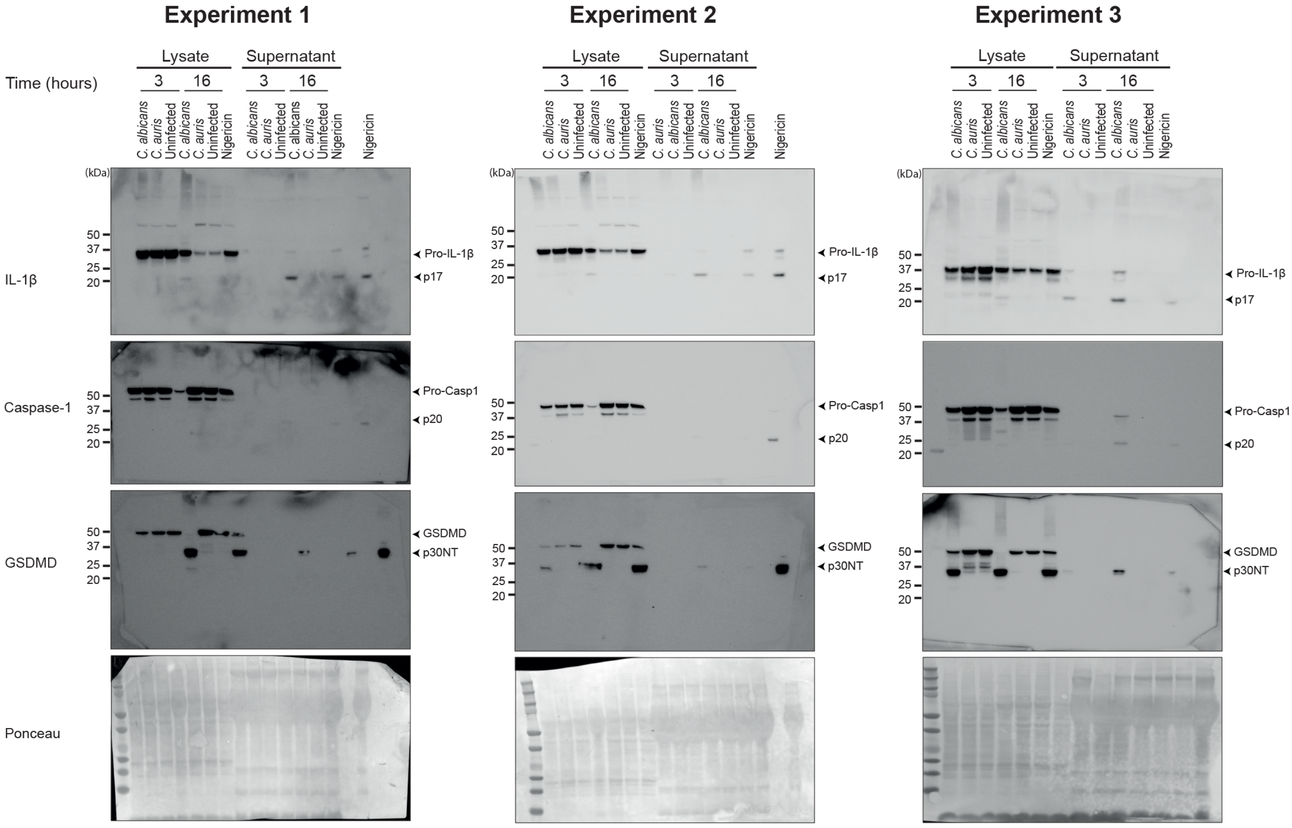
Experiment 1
Experiment 2
Experiment 3
Lysate
Supernatant
Time (hours)
3
16
C. albicans
C. auris
Uninfected
Nigericin
(kDa)
50
37
25
20
IL-1β
Pro-IL-1β
p17
Caspase-1
Pro-Casp1
p20
GSDMD
p30NT
Ponceau

**Supplementary Table 1**

| **Identifier** | **Name** | **Species** | **Genotype** | **Description** | **Source** |
|---|---|---|---|---|---|
| YCAT1118 | 470121 | *C. auris* | Clade I | Clinical isolate | National Mycology Refence Centre, Adelaide. |
| YCAT1200 | B8441 | *C. auris* | Clade I | Clinical isolate, Pakistan | CDC, Atlanta, USA |
| YCAT1191 | B11098 | *C. auris* | Clade I | Clinical isolate, Pakistan | CDC, Atlanta, USA |
| YCAT1195 | B11220 | *C. auris* | Clade II | Clinical isolate, Japan | CDC, Atlanta, USA |
| YCAT1196 | B11221 | *C. auris* | Clade III | Clinical isolate, South Africa | CDC, Atlanta, USA |
| YCAT1198 | B11244 | *C. auris* | Clade IV | Clinical isolate, Venezuela, | CDC, Atlanta, USA |
| YCAT229 | SC5314 | *C. albicans* | Clade I | Clinical isolate | Traven Lab |
| YCAT1336 | 470121 | *C. auris* | *Δtye7::SAT1* | Complete deletion of the *tye7 locus* | This study |
| YCAT1338 | 470121 | *C. auris* | *Δgal4::SAT1* | Complete deletion of the *gal4 locus* | This study |
| YCAT1339 | B8441 | *C. auris* | *Δtye7::SAT1* | Complete deletion of the *tye7 locus* | This study |
| YCAT1341 | B8441 | *C. auris* | *Δgal4::SAT1* | Complete deletion of the *gal4 locus* | This study |
| YCAT1343 | B8441 | *C. auris* | *Δcdc19::SAT1* | Complete deletion of the *cdc19 locus* | This study |

**Supplementary Table 2**

| **Name** | **Purpose** | **5' to 3' sequence** | **Source** |
|---|---|---|---|
| ***Macrophage genes*** | | | |
| *18sRNA (F)* | For qPCR of *18sRNA* | TTGACGGAAGGGCACCACCAG | Tucey *et al*., 2016 |
| *18sRNA (R)* | | GCACCACCACCCACGGAATCG | Tucey *et al*., 2016 |
| *Glut1 (F)* | For qPCR of *Glut1* | AAACTCCAGTCAGCATTTTCATAGG | Tucey *et al*., 2016 |
| *Glut1 (R)* | | CCACGGAGTTGTAAAATATGCTTGT | Tucey *et al*., 2016 |
| *Hk2 (F)* | For qPCR of *Hk2* | TGTTTGTACATTTATTTGCCCTCCT | Tucey *et al*., 2016 |
| *Hk2 (R)* | | GTAGAGCAGAGAACAGGACATTGGT | Tucey *et al*., 2016 |
| *Pfkfb3 (F)* | For qPCR *Pfkfb3* | CTAGCCTACTTCCTCGACAAGAGTG | Tucey *et al*., 2016 |
| *Pfkfb3 (R)* | | ACACTGGCAAAGTGTTGTATTTTGA | Tucey *et al*., 2016 |
| ***Candida genes*** | | | |
| *RDN25 (F)* | For qPCR of *RDN25* | CATATCTAGCAGAAAGCACCGTTC | Tucey *et al*., 2016 |
| *RDN25 (R)* | | CGCGCCGTAGCTTATTCGTATCCAA | This study |
| *CDC19 (F)* | For qPCR of *CDC19* | GCCAAGAAGCCAACTGAGAC | This study |
| *CDC19 (R)* | | GGGTTTCTGGTGACCATCAT | This study |
| *HXK2 (F)* | For qPCR of *HXK2* | TGATCCTTCTCGAGTTGGCTG | This study |
| *HXK2 (R)* | | ATGCCACACACGGACATTCT | This study |
| *PFK1 (F)* | For qPCR of *PFK1* | GTGCAAGGAGGAAACACAGG | This study |
| *PFK1 (R)* | | CCGTGGTGTACACTTTCAGC | This study |
| *PFK2 (F)* | For qPCR of *PFK2* | AGTCTTTGACACCAAAACCGC | This study |
| *PFK2 (R)* | | TGAAACCTCAAGTGGGACGA | This study |
| *TYE7 (F)* | For cloning WT *TYE7* | CATCCTAATGGCACCTCAGCGA | This study |
| *TYE7 (R)* | | GCCGTGCTGGTACTTCCGG | This study |
| *TYE7 inverse (F)* | For deleting *TYE7* ORF | AGAACAAGGCGATGCTGTAC | This study |
| *TYE7 inverse (R)* | | CGCCAGATTAAGTTTACCCACG | This study |
| *GAL4 (F)* | For cloning WT *GAL4* | CGCCTCCGAGGATTACCCA | This study |
| *GAL4 (R)* | | TGCAGTTCTAAATCTGGCCCTAG | This study |
| *GAL4 inverse (F)* | For deleting *GAL4* ORF | CTGAACAGGAGCAATAGG | This study |
| *GAL4 inverse (R)* | | TCGAAGAGTGCTGTGTTG | This study |
| *CDC19 (F)* | For cloning WT *CDC19* | TAACCGCTTGTCAAATCTGTGT | This study |
| *CDC19 (R)* | | GCTCATTTTTCAGCGAGCTTGT | This study |
| *CDC19 inverse (F)* | For deleting *CDC19* ORF | CTAGTGGCTTATGCCGAAAG | This study |
| *CDC19 inverse (R)* | | GTGTAAGATGGGCAATGG | This study |

*F* = Forward primer
*R* = Reverse primer
*ORF* = Open reading frame
